# spike-timing-dependent plasticity rule for single, clustered and distributed dendritic spines

**DOI:** 10.1101/397323

**Authors:** Sabrina Tazerart, Diana E. Mitchell, Soledad Miranda-Rottmann, Roberto Araya

**Affiliations:** Department of Neurosciences, Faculty of Medicine, University of Montreal

**Author notes:** Corresponding author: Roberto Araya Ph.D. Assistant Professor, Department of Neurosciences, Faculty of Medicine, University of Montreal, C.P. 6128 Succursale Centre-Ville, Montreal, Quebec H3C 3J7. These authors contributed equally to this work (Sabrina Tazerart, Diana E. Mitchell).

## Abstract

The structural organization of excitatory inputs that supports spike-timing-dependent plasticity (STDP) remains unknown. Here we performed a spine STDP protocol using two-photon glutamate uncaging (pre) paired with postsynaptic spikes (post). We found that pre-post pairings that trigger LTP (t-LTP) produce shrinkage of the activated spine neck and increase in synaptic strength; and post-pre pairings that trigger LTD (t-LTD) decrease synaptic strength without affecting spine shape. Furthermore, we tested whether this rule could be affected by the activation of two clustered or distributed spines. We show that the induction of t-LTP in clustered spines (<5 µm apart) enhances LTP through a mechanism dependent on spine calcium accumulation and actin polymerization-dependent neck shrinkage, whereas t-LTD was disrupted by the activation of clustered spines but recovered when separated by >40 µm. These results indicate that synaptic cooperativity disrupts t-LTD and extends the temporal window for the induction of t-LTP, leading to STDP only encompassing LTP.

## INTRODUCTION

Dendritic spines, the main recipient of excitatory information in the brain^1^, are tiny protrusions with a small head (∼1 µm in diameter and <1 fL volume) separated from the dendrite by a slender neck. Spines can undergo structural remodeling that is tightly coupled with synaptic function^1, 2, 3, 4^, and are the preferential site for the induction of long-term potentiation (LTP)^4, 5, 6, 7^ and long-term depression (LTD)^8^, thought to be the underlying mechanisms for learning and memory in the brain^9^. A variation of LTP and LTD has been described in pyramidal neurons that involves the pairing of pre- and postsynaptic action potentials, known as spike-timing dependent plasticity (STDP)^10, 11^. In this process, the timing between pre- and postsynaptic action potentials modulates synaptic strength, triggering LTP or LTD^11^. The sign and magnitude of the change in synaptic strength depends on the relative timing between spikes of two connected neurons (the pre- and postsynaptic neuron^12^). The STDP learning rules and their dependency on postsynaptic dendritic depolarization^13, 14^, firing rate^13^, and somatic distance of excitatory inputs^14, 15, 16^ have been extracted from studies using connected neuronal pairs or by using extracellular stimulating electrodes, but the precise location and structural organization of excitatory inputs capable of supporting STDP at its minimal functional unit – the dendritic spine – are unknown.

Activity-dependent spine morphological changes (spine head^4^, neck^2^, or both^17^) have been correlated with changes in synaptic strength in cortical pyramidal neurons by mechanisms involving biochemical and electrical spine changes^1, 6^. Thus, here we asked what patterns of activity and structural organization of excitatory synaptic inputs support the generation of t-LTP and t-LTD, and which morphological, biophysical and molecular changes observed in dendritic spines can account for the induction of t-LTP and t-LTD?

To induce synapse-specific STDP we performed a protocol whereby two-photon (2P) uncaging of a caged glutamate (MNI-glutamate^3^) at a single spine – to mimic synaptic release – is preceded or followed in time (STDP timing window^11^) by a backpropagating action potential (bAP) to trigger t-LTP or t-LTD, respectively. Two-photon uncaging of a caged glutamate at a single spine triggered excitatory postsynaptic potentials (uncaging(u)EPSP) that were recorded in the soma of layer 5 (L5) pyramidal neurons before and after the induction of STDP, while the morphology of the activated spine neck and head was monitored. To induce synapse-specific STDP and monitor calcium levels in the activated spines and parent dendrites we performed a protocol during which we perform nearly simultaneous 2P uncaging of glutamate and 2P calcium imaging of the activated spines and nearby dendrites.

Here, we provide evidence showing that the induction of STDP in single or distributed spines follows a bidirectional Hebbian STDP rule. Furthermore, we show that synaptic cooperativity, induced by the co-activation of only two clustered spines, disrupt t-LTD (< 40 µm distance between spines) and extend the temporal window for the induction of t-LTP (< 5 µm distance between spines) via the generation of differential local calcium signals leading to an STDP rule for clustered inputs only encompassing LTP.

## RESULTS

### Induction of t-LTP in single dendritic spines

To induce t-LTP, we used a repetitive spike-timing protocol (40 times, 0.5 Hz) in which 2P uncaging of glutamate at a single spine was closely followed in time (+7 or +13 ms later, see Methods section) by a bAP (Fig. 1A). We evaluated spine morphology and uEPSP amplitude for 40 min following STDP induction to establish the time course of STDP at individual synapses (Fig. 1C.1 and D.1). In addition, the maximum uEPSP change and concomitant changes in spine morphology observed in each experiment are shown (Fig. 1C.2 and D.2).

**Figure 1:**
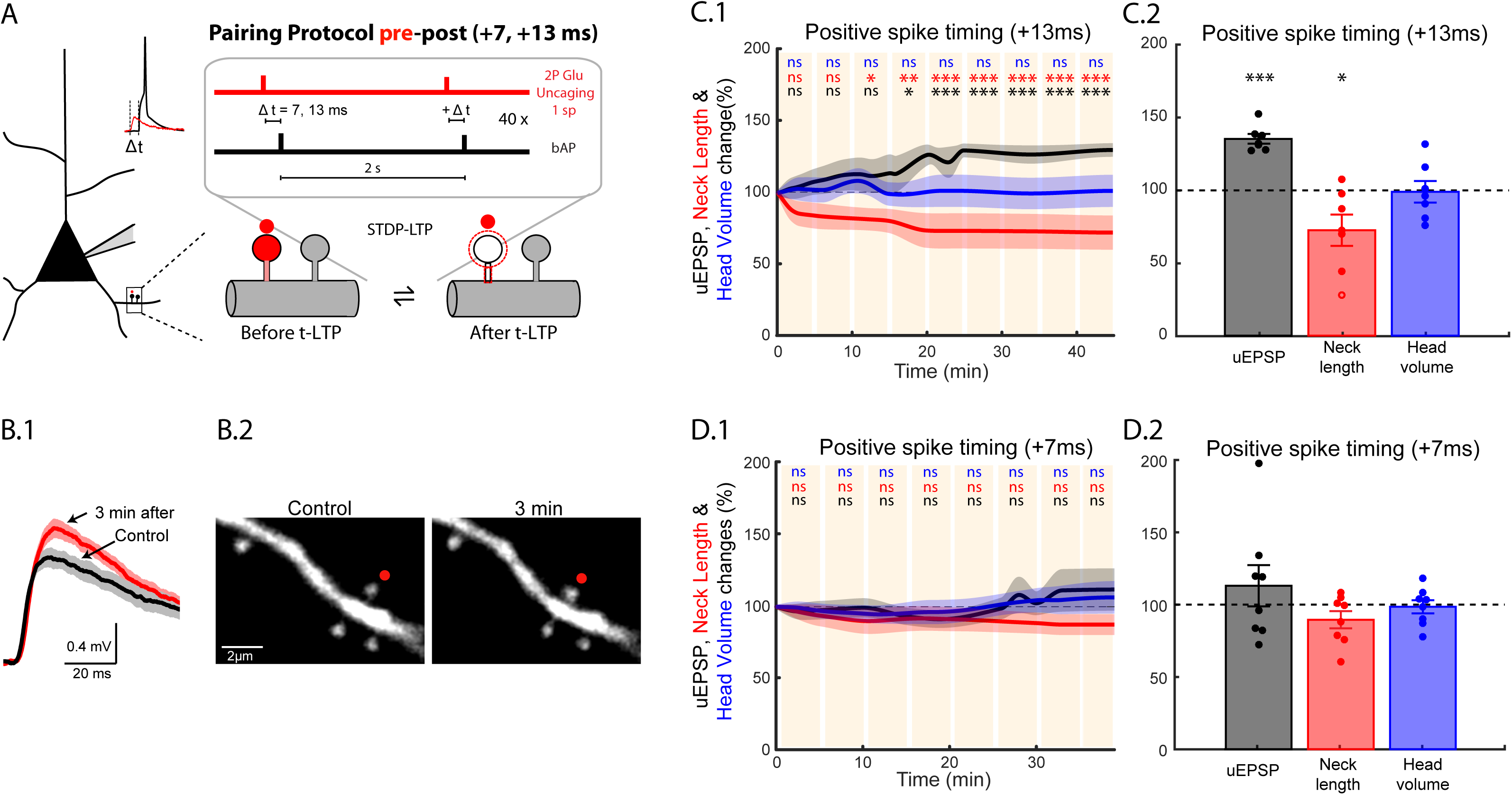
Induction of t-LTP in single dendritic spines. (A) Experimental protocol for the induction of t-LTP with pre-post pairings (two-photon (2P) uncaging of glutamate followed by a bAP) of +7 and +13 ms in single dendritic spines (sp). (B) Example of a t-LTP protocol with pre-post pairing protocol of + 13 ms. (B.1) Average uEPSP response recorded via a whole-cell pipette at the soma of L5 pyramidal neurons before (Control) and after the induction of t-LTP in a selected dendritic spine located in basal dendrites (B.2). Traces in B.1 correspond to an average of 10 depolarizations generated by 2P uncaging over the indicated spine (B.2 red dot) before (black trace) and after the induction of t-LTP (red trace). (C.1) Time course of uEPSP amplitude (black line), neck length (red line) and spine head volume (blue line) changes over the course of 40 min following STDP induction in individual spines at a pre-post timing of + 13 ms. *ns*, not significant; *P < 0.05; **P < 0.01; ***P < 0.001, one-way repeated measures ANOVA followed by post hoc Dunnet’s test. (C.2) Maximum changes in uEPSP amplitude (black bar and dots) and concomitant changes in neck length (red bar and dots) and head volume (blue bar and dots) of the activated spine after the induction of t-LTP at a pre-post timing of +13 ms. *P < 0.05, ***P < 0.001, paired t-test. (D.1) Time course of uEPSP amplitude (black line), neck length (red line) and spine head volume (blue line) changes over the course of ∼40 min following STDP induction in individual spines at a pre-post timing of + 7 ms. *ns*, not significant; *P < 0.05, one-way repeated measures ANOVA followed by post hoc Dunnet’s test. (D.2) Maximum changes in uEPSP amplitude (black bar and dots) and corresponding changes in neck length (red bar and dots) and head volume (blue bar and dots) of the activated spine after the induction of t-LTP at a pre-post timing of +7 ms. Shaded light area in C.1 and D.1, and error bars in C.2 and D.2 represent ±SEM. Time 0 in C.1 and D.1 is the time when the 40 pre-post repetitions of the induction protocol were completed.

A repetitive pre-post pairing protocol of +13 ms reliably induced t-LTP (significant increase in the uEPSP amplitude over time, P < 0.001, n = 7 experiments, from 6 neurons, from 6 mice, Fig. 1B.1 and C.1), and shortening of the activated spine neck within a few minutes (P < 0.001), with no significant change in spine head volume (n = 7, Fig. 1B.2 and C.1). These results were also consistent when we considered the maximum uEPSP change in amplitude from each experiment and concomitant changes in spine morphology (uEPSP = 134.21 ± 3.29%, P < 0.001, n = 7; neck length = 71.88 ± 10.66%, P < 0.05, n = 7; spine head volume = 98.11 ± 7.34%, P = 0.81, n = 7) (Fig. 1C.2). We obtained similar results when we instead considered the average of all the values obtained following t-LTP induction for uEPSP amplitude, neck length and head volume (Fig. S1). This effect was specific to the activated spine (Fig. 1B.2), with neighbouring spines having no appreciable changes in their neck length or head volume (neck length = 98.09 ± 5.06%, P = 0.71, n = 13; head volume = 103.01 ± 3.61%, P = 0.42, n = 14). Control experiments showed that there was no significant change in uEPSP amplitude or spine morphology following the STDP protocol when either action potentials or synaptic stimulation were applied in isolation, as well as when we monitored the long-term stability of these parameters without any STDP protocol (Fig. S2).

A pre-post pairing of + 7 ms showed a non-significant tendency for the induction of LTP following t-LTP induction, and a non-significant tendency for the shrinkage of the activated spine neck (n = 8 experiments, from 8 neurons, from 8 mice, Fig. 1D.1). When the maximum change in uEPSP amplitude from each experiment and the concomitant changes in spine morphology were analyzed, we saw no significant changes in uEPSP amplitude and spine morphology (Fig. 1D.2, uEPSP = 112.27 ± 14.19%, P = 0.42, n = 8; neck length = 88.90 ± 5.89%, P = 0.10, n = 8; head volume = 97.81 ± 4.54%, P = 0.64, n = 8) (Fig. 2E). Similar results were observed when we instead considered the average of all the values obtained following t-LTP induction (+7 ms) for uEPSP amplitude, neck length and head volume (Fig. S1). Because voltage has been shown to be an important factor in the induction of t-LTP and t-LTD, we verified that the initial uEPSP amplitudes were not significantly different for pre-post pairing protocols of +13 ms versus +7 ms (uEPSP: 0.62 ± 0.14 versus 0.53 ± 0.16 mV, P = 0.68; Fig. S3).

**Figure 2:**
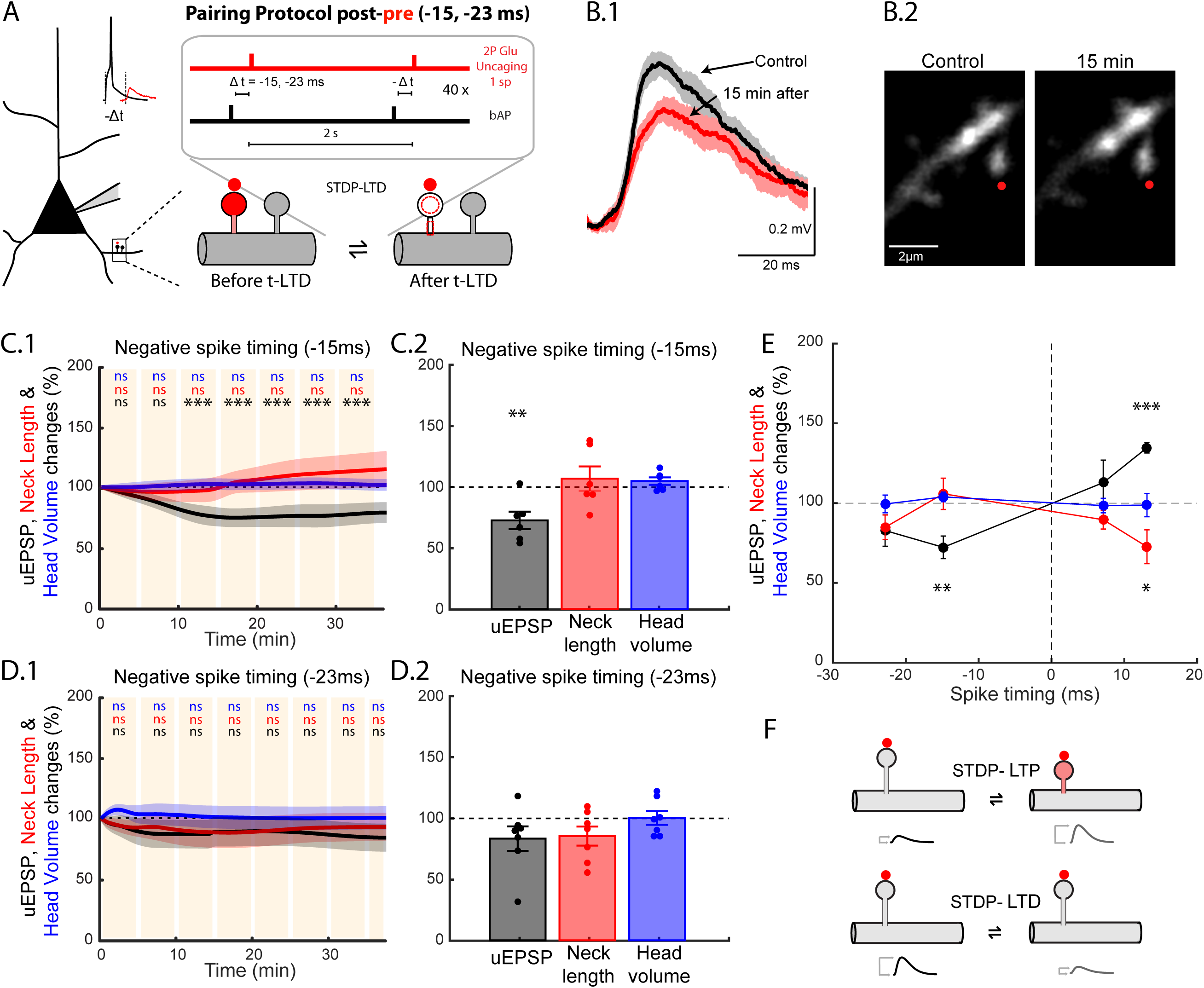
Induction of t-LTD in single dendritic spines. (A) Experimental protocol for the induction of t-LTD with post-pre pairings protocols of −15 and −23 ms in single dendritic spines. (B) Example of a t-LTD protocol with post-pre pairings protocol of −15 ms. (B.1) Average uEPSP response recorded via a whole-cell pipette at the soma of L5 pyramidal neurons before (Control) and after the induction of t-LTD in selected dendritic spines located in basal dendrites (B.2). Traces in B.1 correspond to an average of 10 uEPSP generated by the 2P uncaging of glutamate at the indicated spine (B.2, red dot) before (black trace) and after the induction of t-LTD (red trace). (C.1) Time course of uEPSP amplitude (black line), neck length (red line) and spine head volume (blue line) changes over the course of ∼35 min following STDP induction in individual spines at a post-pre timing of −15 ms. *ns*, not significant; ***P < 0.001, one-way repeated measures ANOVA followed by post hoc Dunnet’s test. (C.2) Maximum changes in uEPSP amplitude (black bar and dots) and concomitant changes in neck length (red bar and dots) and head size of the activated spine (blue bar and dots) after the induction of t-LTD at a post-pre timing of −15 ms. **P < 0.01, paired t test. (D.1) Time course of uEPSP amplitude (black line), neck length (red line) and spine head volume (blue line) changes over the course of ∼ 40 min following STDP induction in individual spines at a post-pre timing of −23 ms. *ns*, not significant, one-way repeated measures ANOVA followed by post hoc Dunnet’s test. (D.2) Maximum changes in uEPSP amplitude (black bar and dots) and corresponding changes in neck length (red bar and dots) and head volume (blue bar and dots) of the activated spine after the induction of t-LTD at a post-pre timing of −23 ms. Shaded light area in C.1 and D.1, and error bars in C.2 and D.2 represent ±SEM. Time 0 in C.1 and D.1 is the time when the 40 pre-post repetitions of the induction protocol were completed. (E) STDP learning rule for single dendritic spines: Plot illustrating the maximum changes in uEPSP amplitude (black data points), and concomitant changes in neck length (red data points) and head volume (blue data points) after the induction of t-LTP (temporal offset +13 ms and +7 ms) and t-LTD (temporal offset −15 ms and −23 ms). *P < 0.05, **P < 0.01, paired t test. (F) Diagram indicating the uEPSP amplitude and spine neck morphological changes observed after the induction of t-LTP and t-LTD.

These results indicate that for the pairing times tested, there is a preferred pre-post t-LTP pairing time-window (+ 13 ms) at which activated spines in basal dendrites from L5 pyramidal neurons undergo a significant increase in synaptic strength, and a concomitant neck shrinkage (Fig. 2E).

### Induction of t-LTD in single dendritic spines

We then studied t-LTD in single spines by using a repetitive spike-timing protocol (40 times, 0.5 Hz) in which 2P uncaging of glutamate at a single spine was preceded in time (−15 or −23 ms) by a bAP (post-pre protocol, Fig. 2A). When postsynaptic spikes preceded presynaptic firing by 15 ms (i.e., −15 ms), a significant reduction of the uEPSP amplitude occurred within a few minutes following induction of t-LTD (n= 6 experiments, from 5 neurons and 5 mice, Fig. 2B.1 and C.1, P < 0.001), with no significant changes in spine neck length or head dimension (Fig. 2B.2 and C.1). Furthermore, when the maximum change in uEPSP amplitude after the induction of t-LTD in single spines at pairings of −15 ms was analyzed (Fig. 2C.2) we also observed a significant depression of uEPSP amplitude (uEPSP = 71.52 ± 7.07%, P < 0.01, n = 6), with no significant changes in spine morphology (Fig. 2B.2 and C.2, neck length = 105.54 ± 9.85%, P = 0.62, n = 6; head volume = 103.25 ± 3.02%, P = 0.33, n = 6). Interestingly, after the induction of t-LTD in single spines when postsynaptic spikes preceded presynaptic firing by 23 ms (i.e., −23ms) there were no significant changes in the amplitude of the uEPSPs or in the spine neck length and head dimensions for the duration of the recordings (n= 7 experiments, from 7 neurons and 7 mice, Fig. 2D.1). These results were also consistent with analyses of the maximal uEPSP change in each experiment and concomitant spine morphology (Fig. 2D.2: uEPSP = 82.09 ± 9.89%, P = 0.12, n = 7; neck length = 84.15 ± 7.73%, P = 0.09, n = 7; head volume = 98.81 ± 5.57%, P = 0.84, n = 7). There was no significant difference between the initial uEPSP amplitude for post-pre pairing protocols of −15 ms versus −23 ms (EPSP: 0.59 ± 0.07 versus 0.49 ± 0.08 mV, respectively, P = 0.42; Fig. S3). We obtained similar results when we instead considered the average of all the values obtained following t-LTD induction in single spines for uEPSP amplitude, neck length and head volume (Fig. S1). This indicates that for the pairing times tested, only a post-pre t-LTD pairing time-window of −15 ms can effectively induce LTD in single dendritic spines in the basal dendrites from L5 pyramidal neurons.

Taken together these results show that the induction of t-LTP and t-LTD in single spines follows a Hebbian-STDP learning rule that is bidirectional, and favors presynaptic inputs that precede postsynaptic spikes and depresses presynaptic inputs that are uncorrelated with postsynaptic spikes at a very precise and narrow temporal window (+13 ms for the generation of t-LTP and - 15 ms for t-LTD, Fig. 2E-F). The single spine STDP rule we observed has a narrower post-pre LTD induction pairing time window than previously observed in connected pairs of L2/3^15, 18^ and L5 pyramidal neurons^13^ – where the presynaptic control of t-LTD via an mGluR and/or cannabinoid type 1 receptor-dependent mechanism^19, 20, 21, 22^ could plausibly account for these differences.

### Induction of t-LTP in clustered dendritic spines

It has been suggested that STDP not only depends on spike timing and firing rate but also on synaptic cooperativity and the amount of voltage generated at the postsynaptic site^13, 14^. Furthermore, it has been shown in mouse hippocampal CA1 pyramidal cells that the induction of t-LTP in a single spine in low extracellular Mg^2+^ conditions can reduce the threshold for the induction of plasticity of a neighboring spine (separated by < 10 µm) activated 90 seconds later^7^. However, a direct demonstration of synaptic cooperativity for synchronous, or nearly synchronous synaptic inputs, in physiological extracellular Mg^2+^ conditions, and at the level of single spines for the induction of STDP in the dendrites of pyramidal neurons has yet to be obtained. Hence, we next directly tested if synaptic cooperativity, marked by the co-induction of t-LTP in clustered dendritic spines from basal dendrites of L5 pyramidal neurons, and the local spatio-temporal summation of inputs, can generate a local dendritic depolarization and local calcium signals, which are high enough to disrupt the single spine STDP learning rule described in Fig. 2E. To test this, a two spine STDP protocol (forty 2P uncaging pulses, pulse duration 2ms, 0.5Hz, see Methods section) was performed, whereby nearly simultaneous 2P uncaging of caged glutamate in clustered (distance between spines < 5 µm) spines was followed in time by a bAP to trigger t-LTP (Fig. 3A). With this protocol, we investigated whether activating clustered spines extended the pre-post timing window capable of generating LTP by increasing the degree of depolarization immediately before the postsynaptic spike at timings where plasticity was not reliably generated. Specifically, we induced t-LTP in two clustered spines at pre-post timings of +7 ms, and surprisingly found that this protocol was in fact capable of effectively and significantly generating increases in uEPSP amplitude (Fig. 3B.1) and the concomitant shrinkage of the activated spine neck, with no apparent changes in its spine head size (Fig. 3B.2). Pooled data showed that significant increases in uEPSP amplitude and shrinkage of the spine neck of the activated spines occurs only a few minutes post t-LTP induction and lasted for the duration of the recordings (Fig. 3C, P < 0.01, n = 10 spines from 8 experiments, 6 neurons and 6 mice), with no significant changes in the spine head size (Fig. 3C, n = 16 spines from 8 experiments, 6 neurons, 6 mice). Similar results were observed when we analyzed the maximal change in uEPSP amplitude in each experiment and the concomitant spine neck length and head size of the two clustered spines after induction of t-LTP at pairings of + 7 ms (Fig. 3H; uEPSP = 130.86 ± 8.18%, P < 0.01, n = 8; neck length = 73.22 ± 5.84%, P < 0.01, n = 10; head volume =102.00 ±2.58%, P = 0.45, n = 16). We obtained similar results when we instead considered the average of all the values obtained following t-LTP induction in two clustered spines for uEPSP amplitude, neck length and head volume (Fig. S1). In control experiments, no significant change in uEPSP amplitude or spine morphology was observed when we monitored the long-term stability of these parameters without any STDP protocol (Fig. S2B). These results indicate that synaptic cooperativity of nearly simultaneous synaptic inputs – shown by the induction of t-LTP in only two clustered spines (< 5 µm apart) – is sufficient to significantly trigger synaptic potentiation and shrinkage of the activated spine necks at a pre-post timing that is otherwise ineffective at generating significant morphological changes and synaptic potentiation when only one spine is being activated (for comparison between one versus two clustered spines see Fig. S4). Hence, the synaptic cooperativity of only two neighbouring synaptic inputs onto spines (< 5 µm apart) in the basal dendrites of L5 pyramidal neurons extends the pre-post timing window that can trigger potentiation (Fig. 3H, and compare Fig. 3C with Fig. 1D.1).

**Figure 3:**
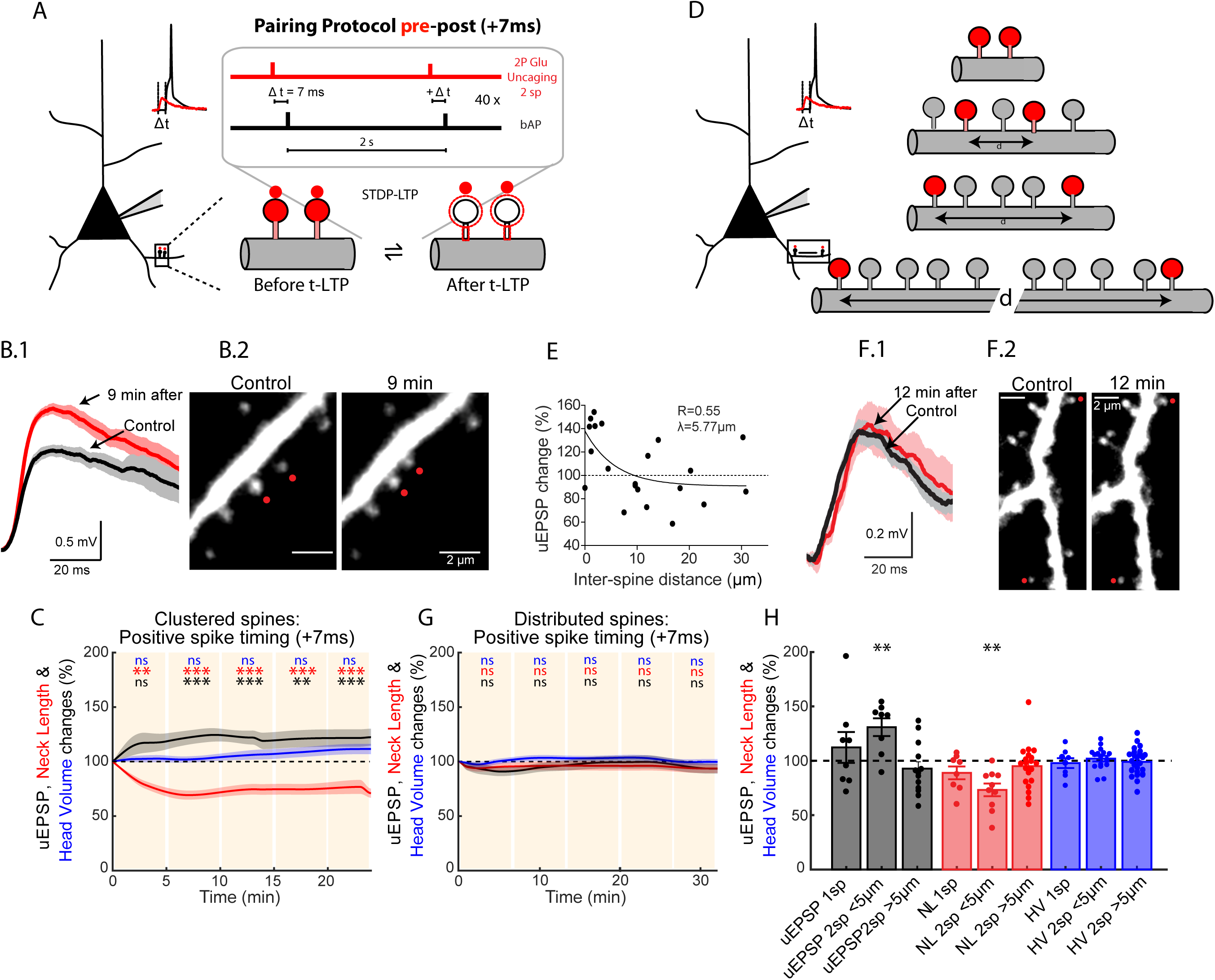
Induction of t-LTP in clustered and distributed dendritic spines. (A) Experimental protocol for the induction of t-LTP (pre-post timing of + 7 ms) in two clustered dendritic spines (< 5 µm distance between spines). (B) Example of one experiment where two neighbouring dendritic spines were activated with a pre-post t-LTP protocol. (B.1) Average uEPSP response recorded via a whole-cell pipette at the soma of L5 pyramidal neurons before (Control) and after the induction of t-LTP in two selected dendritic spines located in a basal dendrite (red dots, B.2). Traces in B.1 correspond to an average of 10 uEPSP generated by the 2P uncaging at the indicated spines (B.2) before (black trace) and after the induction of t-LTP (red trace). (C) Time course of uEPSP amplitude (black line), neck length (red line) and spine head volume (blue line) changes over the course of ∼25 min following STDP induction in clustered spines (< 5µm distance between spines) at a pre-post timing of + 7 ms. *ns*, not significant; **P < 0.01, ***P < 0.001, one-way repeated measures ANOVA followed by post hoc Dunnet’s test. (D) Experimental protocol for the induction of t-LTP (pre-post timing of + 7 ms) in two distributed spines (> 5µm distance between spines). (E) Induction of t-LTP (increase in uEPSP amplitude, more than 100%) in two nearly simultaneously activated spines at a pre-post timing of + 7 ms is dependent on inter-spine distance. Data points represent mean. We use the extracted value of λ as a boundary between clustered and distributed spines. (F) Example of one experiment where two distributed dendritic spines (> 5µm distance between spines) were activated with a pre-post timing of + 7ms. (F.1) Average uEPSP response recorded via a whole-cell pipette at the soma of L5 pyramidal neurons before (Control) and after the induction of t-LTP in two selected dendritic spines located in a basal dendrite (red dots, F.2). Traces in F.1 correspond to an average of 10 uEPSP generated by the 2P uncaging at the indicated spines (F.2) before (black trace) and after the induction of t-LTP (red trace). (G) Time course of uEPSP amplitude (black line), neck length (red line) and spine head volume (blue line) changes over the course of ∼30 min following STDP induction in distributed spines (> 5µm distance between spines) at a pre-post timing of + 7 ms. *ns*, not significant; **P < 0.01, ***P < 0.001, one-way repeated measures ANOVA followed by post hoc Dunnet’s test. (H) Pooled data of the maximum changes in uEPSP amplitude (black bar and dots) and concomitant changes in neck length (red bar and dots) and head volume (blue bar and dots) of individual (1sp), clustered (< 5µm) and distributed (> 5µm) spines (2sp) after the induction of t-LTP at a pre-post timing of +7ms. **P < 0.01, paired t test. Shaded light area in B.1, F.1C and G and error bars in H represent ±SEM. Time 0 in C and G is the time when the 40 pre-post repetitions of the induction protocol are completed. NL = neck length, HV = head volume.

### Induction of t-LTP in distributed dendritic spines

To precisely study the effect of inter-spine distance of the nearly simultaneously activated spines on synaptic cooperativity and the induction of t-LTP at pairings of +7 ms, we performed experiments where the activated spines were further away (> 5 µm apart) from each other. When we correlated 1) the inter-spine distance and the time at which the maximal uEPSP change was observed for each experiment, or 2) the mean change - obtained when we analyzed the average change in uEPSP from all the times tested for each experiment (Fig. S5A) - following t-LTP induction, we found that as two activated spines were further away from each other, the less t-LTP was induced (Fig. 3E-F, Fig. S5A). When the maximal uEPSP change per experiment was analyzed, we observed that the induction of t-LTP decayed exponentially as a function of inter-spine distance with a length constant (λ) of 5.7 µm (Fig. 3E). This value of λ was similar to that obtained when we analyzed the average uEPSP change of all time points tested for each experiment after t-LTP induction (Fig. S5A, P= 0.87, obtained when comparing the maximal versus the average change in uEPSP used to calculate λ). Therefore we used this value as the boundary between clustered (< 5 µm) and distributed (> 5 µm) spines for the induction of t-LTP at pairings of + 7ms. Clustered spines were located in the same basal dendrite (n = 8/8 pairs), while distributed spines were either in sister branches emanating from the same bifurcation point (n = 7/13 pairs) or in the same basal dendrite (n = 6/13 pairs). By separating the data in this manner, the t-LTP protocol (+ 7ms) in distributed spines (> 5 µm, Fig. 3D) failed to induce t-LTP (Fig. 3F.1) or changes in spine head size and neck length at all the times tested post t-LTP induction (Fig. 3G, n = 13 experiments, n = 26 spines, from 11 neurons and 7 mice). For comparisons between the activation of two clustered versus two distributed spines with a pre-post timing of + 7ms see Fig. S4. When we analyzed the maximum uEPSP change in amplitude in each experiment and the concomitant changes in spine head volume and neck length in the activated spines after t-LTP induction in two distributed spines, we also observed a complete loss of t-LTP induction, with no changes in spine morphology (Fig. 3H; uEPSP = 92.52 ± 6.570%, P = 0.28, n = 13 experiments; neck length = 94.76 ± 4.51%, P = 0.26, n = 20 spines; head volume = 98.65 ± 2.41%, P = 0.58, n = 26 spines, from 13 experiments performed in 11 neurons from 7 mice). Similar results were obtained when we instead considered the average of all the values obtained following t-LTP induction (+7ms) in distributed spines for uEPSP amplitude, neck length and head volume (Fig. S1). In summary, this data shows that the induction of t-LTP in nearly synchronously activated spines at pre-post timings of + 7 ms is disrupted when the two activated spines were more than 5 µm apart in the basal dendrites of L5 pyramidal neurons. These results uncover a two-spine activation crosstalk-spatial barrier of 5 µm for the induction of t-LTP to occur at pre-post timings of + 7 ms.

### Molecular mechanisms responsible for the generation of t-LTP in dendritic spines

The results led us to consider the possible mechanisms underlying the generation of t-LTP at the level of single spines. Specifically, we asked why the induction of t-LTP in single and clustered spines is associated with the shrinkage of the activated spine neck. We and others have reported that the induction of LTP can trigger activity dependent changes in neck length^2, 17^ and spine head size^4, 6, 17^, and that the amplitude of uEPSP recorded at the cell soma is inversely proportional to the length of the activated spine neck^2, 23, 24^. However the mechanisms by which the t-LTP-induced neck shrinkage is associated with synaptic plasticity remains unknown. Numerical simulations show that the EPSP amplitude/neck length correlation can be explained by variations in synaptic conductance, electrical attenuation through the neck, or a combination of the two^2^. Nevertheless, solutions that rely exclusively on the passive electrical attenuation of synaptic inputs through the spine neck assume very high (> 2 GOhm) neck resistance^2^, which are at odds with recent spine neck resistance estimations^25, 26^. These results suggest that the control of AMPA receptor content in spines could contribute significantly to the observed t-LTP-dependent changes in synaptic strength. To experimentally study the contribution of AMPA receptors to these phenomena, we performed t-LTP experiments in two clustered spines from L5 pyramidal neurons recorded via patch pipettes loaded with intracellular solution containing 200 µM PEP1-TGL – a peptide that inhibits AMPA receptor incorporation to the postsynaptic density (PSD) by blocking GluR1 C-terminus interaction with PDZ domains at the PSD^27^ (Fig. 4A). PEP1-TGL incubation by itself did not trigger a run-down of uEPSP amplitude or changes in spine morphology over time (Fig. S6A). Pooled data from experiments where a repetitive pre-post pairing protocol of + 7 ms was used to activate clustered spines in the presence of PEP1-TGL show that the peptide completely inhibited t-LTP for the duration of the experiment (Fig. 4B.1 and C.1), but had no effect on the t-LTP-induced shrinkage of the activated spine necks (Fig. 4B.2 and C.1, P < 0.001, n = 6 spines, from 5 experiments, 5 neurons and 5 mice) or in modifying spine head size (n = 10 spines, from 5 neurons and 5 mice, Fig. 4C.1). Furthermore, when we analyzed the maximum change in uEPSP amplitude and the concomitant spine morphology after the induction of t-LTP in the presence of PEP1-TGL, we also found an inhibition of t-LTP, but no effect on the t-LTP-induced shrinkage of the spine neck (Fig. 4B.2- C.2, uEPSP = 94.82 ± 14.82%, P = 0.74, n = 5 experiments, from 5 neurons and 5 mice; neck length = 83.92 ± 5.35%, P < 0.05, n = 6; head volume = 100.08 ± 3.23%, P = 0.98, n = 10). We obtained similar results when we instead considered the average of all the values obtained following t-LTP induction in two clustered spines in the presence of PEP1-TGL for uEPSP amplitude, neck length and head volume (Fig. S1). No significant difference was observed between the initial uEPSP amplitude for pre-post pairing protocols of +7 ms with versus without PEP1-TGL (uEPSP: 1.06 ± 0.2 versus 1.16 ± 0.28 mV, P = 0.81; Fig. S3). These results indicate that GluR1 receptor incorporation into the PSD - via its interaction with PDZ domains - is required for the induction of t-LTP in spines. However, the role of the spine neck shrinkage in AMPA receptor incorporation into the PSD and ultimately on the induction of t-LTP remains open.

**Figure 4:**
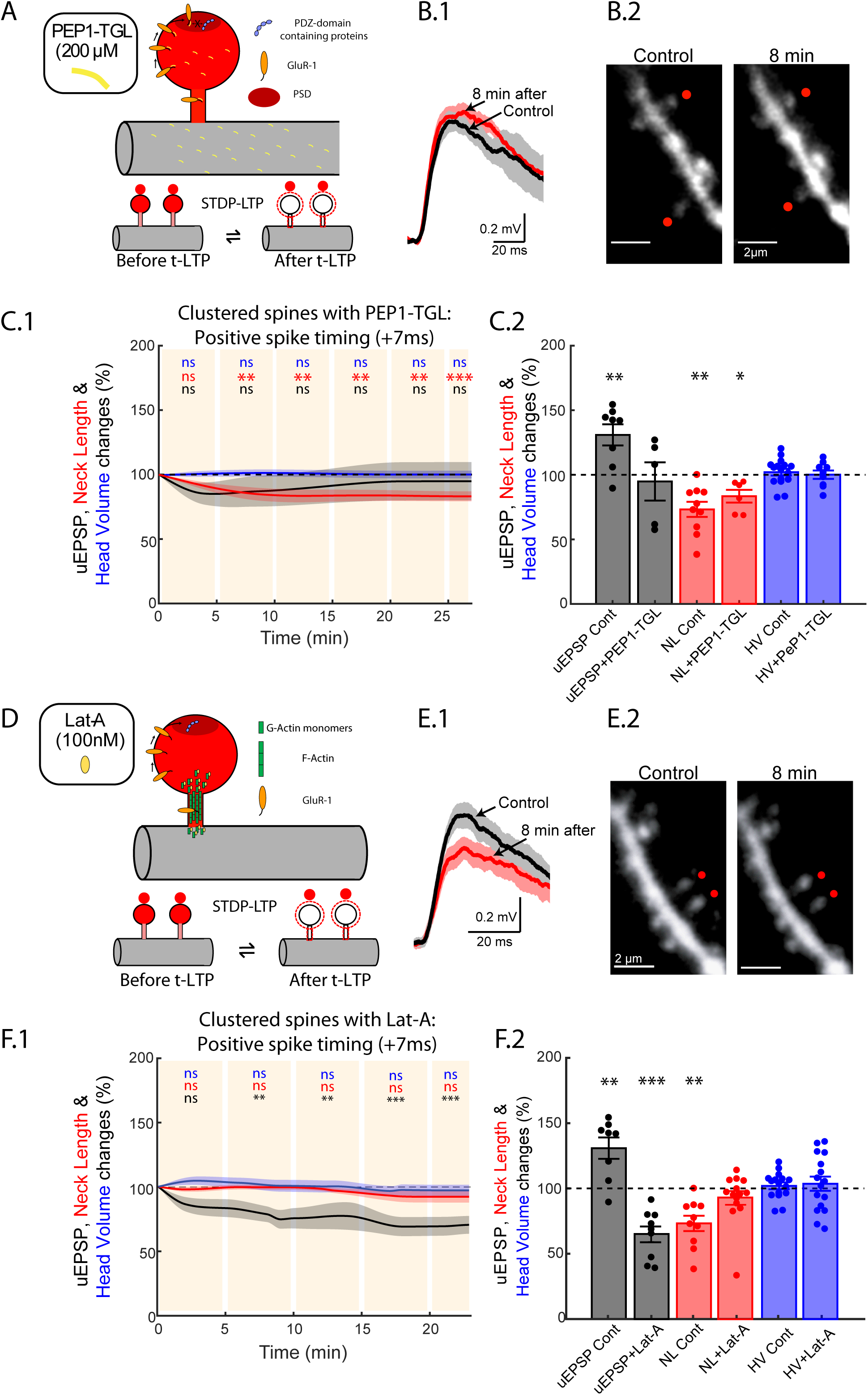
Molecular mechanisms responsible for the induction of t-LTP. (A) Experimental protocol for the induction of t-LTP (pre-post timing of + 7 ms) in two clustered dendritic spines (< 5 µm) in the presence of PEP1-TGL (200 µM). (B) An example of one experiment where two neighbouring dendritic spines were activated with a pre-post t-LTP protocol in the presence of PEP1-TGL. (B.1) Average uEPSP recorded via a whole-cell pipette at the soma of L5 pyramidal neurons before (Control) and after the induction of t-LTP in two selected dendritic spines located in basal dendrites (B.2). Traces in B.1 correspond to an average of 10 uEPSP generated by the 2P uncaging of glutamate at the two indicated spines (red dots, B.2) before (black trace) and after the induction of t-LTP in the presence of PEP1-TGL (red traces). (C.1) Time course of uEPSP amplitude (black line), neck length (red line) and spine head volume (blue line) changes over the course of ∼28 min following STDP induction in clustered spines at a pre-post timing of + 7 ms in the presence of PEP1-TGL. *ns*, not significant; **P < 0.01, ***P < 0.001, one-way repeated measures ANOVA followed by post hoc Dunnet’s test. (C.2) Pooled data of the maximum changes in uEPSP amplitude (black bar and dots) and concomitant changes in neck length (red bar and dots) and head volume (blue bar and dots) of the activated clustered spines after the induction of t-LTP at a pre-post timing of +7ms in control conditions (Cont) and in the presence of PEP1-TGL (PEP1-TGL). *P < 0.05, **P < 0.01, paired t test. Time 0 is the time when the 40 pre-post repetitions of the induction protocol were completed. (D) Experimental protocol for the induction of t-LTP (pre-post timing of + 7 ms) in two clustered dendritic spines (< 30µm) in the presence Latrunculin-A (Lat-A, 100nM). (E) Example of one experiment where two neighbouring dendritic spines were activated with a pre-post t-LTP protocol in the presence of Lat-A. (E.1) Average uEPSP recorded via a whole-cell pipette at the soma of L5 pyramidal neurons before (Control) and after the induction of t-LTP in two selected dendritic spines (red dots, E.2) located in basal dendrites (E.2). Traces in E.1 correspond to an average of 10 uEPSP generated by the 2P uncaging over the two indicated spines (red dots, E.2) before (black trace) and after the induction of t-LTP in the presence of Lat-A (red trace). (F.1) Time course of uEPSP amplitude (black line), neck length (red line) and spine head volume (blue line) changes over the course of ∼12 min following STDP induction in clustered spines at a pre-post timing of + 7 ms in the presence of Lat-A. *ns*, not significant; **P < 0.01, ***P < 0.001, one-way repeated measures ANOVA followed by post hoc Dunnet’s test. (F.2) Pooled data of the maximum changes in uEPSP amplitude (black bar and dots) and concomitant changes in neck length (red bar and dots) and head volume (blue bar and dots) of the activated clustered spines after the induction of t-LTP at a pre-post timing of +7 ms in control conditions (Cont) and in the presence of Lat-A (Lat-A). **P < 0.01, ***P < 0.001, paired t test. NL = neck length, HV = head volume. Shaded light area in C.1 and F.1 and error bars in C.2 and F.2 represent ±SEM. Time 0 is the time when 40 pre-post repetitions of the induction protocol were completed.

Experimental and theoretical studies have indicated that lateral diffusion of AMPA receptors into and out of the spine head can be restricted by the spine neck geometry^28, 29, 30, 31^. In particular, lateral diffusion of AMPA receptors into and out of mushroom spines (long-necked spines) has been shown to be significantly slower than that observed in stubby spines (small-necked spines)^28^ – which is supported by studies that show reduced diffusion of membrane proteins located in spine necks^32^. In addition, quantitative models using realistic spine morphologies indicate that decreasing the radius and increasing the spine neck length increases the retention of AMPA receptors at the synapse^30^, even when their interaction with scaffolding cytoskeletal proteins is neglected^31^. Actin is highly enriched in the spine neck and head^33^, and plays an important role in anchoring AMPA receptors in the spine^34^ and AMPA receptor trafficking^35^, being instrumental for synaptic transmission and plasticity^36, 37, 38^. Hence, to address the role that t-LTP-induced neck shrinkage has on AMPA receptor lateral trafficking to the PSD, and the generation of t-LTP in the activated spines we focused on actin dynamics. We used the actin polymerization inhibitor latrunculin A (Lat-A)^34, 36, 38^ (Fig. 4D), which did not trigger any run-down of uEPSP amplitude or changes in spine morphology over time in the absence of STDP induction (Fig. S6B), to test the potential role of actin dynamics on the spine induction of t-LTP, and on the neck shrinkage and AMPA receptor incorporation into the PSD in the activated spines (Fig. 4D). The induction of t-LTP at pre-post pairings of +7 ms in two clustered spines in the presence of 100 nM Lat-A completely blocked the shrinkage of the activated spine necks and the induction of t-LTP (Fig. 4E and F.1, n = 9 spines, from 8 neurons and 5 mice), inducing instead a significant reduction in uEPSP amplitude over time (Fig. 4F.1, P < 0.001, n = 9 experiments, from 8 neurons and 5 mice). These observations were also consistent with analyses of the maximal change in uEPSP amplitude in each experiment and concomitant spine morphology post t-LTP induction in the presence of Lat-A (Fig. 4F.2, uEPSP = 65.25 ± 6.22%, P < 0.001, n = 9 experiments; neck length = 93.07 ± 5.15%, P = 0.2, n = 14 spines; head volume = 103.6 ± 5.39%, P = 0.51, n = 16 spines).

We obtained similar results when we instead considered the average of all the values obtained following t-LTP induction in two clustered spines in the presence of 100 nM Lat-A for uEPSP amplitude, neck length and head volume (Fig. S1).

No significant difference was observed between initial uEPSP amplitudes for pre-post pairing protocols of +7 ms with versus without Lat-A (uEPSP: 0.66± 0.1 versus 1.16 ± 0.28 mV, P = 0.11; Fig. S3). The lack of run-down of uEPSP amplitude over time in neurons treated with Lat-A in the absence of STDP induction (Fig. S6B), but the significant depression in uEPSP after the induction of t-LTP suggests that the induction of plasticity, and the rearrangement of actin filaments de-stabilized AMPA receptors, leading to removal from the PSD.

In summary, these results show that actin polymerization is required for the t-LTP-dependent neck shrinkage and the induction of plasticity. Our findings further suggest that the induction of t-LTP occurs via a mechanism that involves a neck-shrinkage-dependent facilitated diffusion of GluR1 subunits from the spine neck to the head, and subsequent incorporation into the PSD. We hypothesize that a shorter and wider neck facilitates the transport of AMPA receptors into the spine head (Fig. 4D), a mechanism that is required for the induction of t-LTP.

### Induction of t-LTD in clustered and distributed dendritic spines

We then studied whether the induction t-LTD in single spines observed at pairings of −15 ms could be affected by synaptic cooperativity. Our reasoning was based on two previous observations which suggest that 1) t-LTP induction in the distal dendrites of L5 pyramidal neurons (layer 3-L5 pyramidal neuron pairs) triggers LTD instead of LTP, and 2) that LTD can be converted into LTP by increasing the local voltage^14^. We hypothesized that the induction of t-LTD in single spines depends on the degree of local depolarization and hence, LTD can be disrupted by the activation of neighboring spines. To test this, we performed repetitive spike-timing protocol (40 times, 0.5 Hz) in which 2P uncaging of glutamate at two spines (separated by up to 100 µm) was preceded in time (−15 ms) by a bAP (Fig. 5A.2 and Fig. S7A). Surprisingly, we found that this t-LTD protocol failed to induce any change in uEPSP amplitude or spine head volume with only a slight but significant reduction in spine neck length (Fig. S7A). When we analyzed the maximum uEPSP change in amplitude in each experiment and the concomitant morphological alterations in the activated spines after t-LTD induction, we observed a complete inhibition in the induction of t-LTD, and no change in spine morphology (Fig. S7B; uEPSP = 93.22 ± 6.29%, P = 0.30, n = 17 experiments from 14 neurons and 14 mice; neck length = 88.56 ± 5.69%, P = 0.06, n = 23 spines; head volume = 102.41 ± 6.10%, P = 0.69, n = 34 spines). To more precisely characterize the effect of activating two spines on the induction of t-LTD, we correlated the inter-spine distance and either 1) the maximum uEPSP change from each experiment or 2) the mean uEPSP change of all the times tested from each experiment following STDP induction (see Methods). We found that as the two activated spines were further away from each other, the more t-LTD was recovered (Fig. 5D, each data point corresponds to the time at which the maximal change in uEPSP was observed after t-LTD induction). Specifically, the uEPSP change decayed exponentially as a function of inter-spine distance with a length constant (λ) of 43.5 µm. This value of λ was similar to that obtained when we analyzed the average uEPSP change of all time points tested for each experiment after t-LTD induction (Fig. S5B, P= 0.8, obtained by comparing the maximal versus the average change in uEPSP used to calculate λ).Therefore, we used this value as a boundary between clustered (< 40 µm) and distributed (> 40 µm) spines. Using this classification, clustered spines were located in the same dendrite (n = 11/12 pairs) or in sister branches emanating from the same bifurcation point (n = 1/12 pairs), while distributed spines were always located on separate dendrites (n = 5/5 pairs). When we separated our data in this manner, the t-LTD protocol in two clustered spines (Fig. 5A) failed to induce t-LTD (Fig. 5B.1) or changes in spine head size at all the times tested post t-LTD induction (Fig. 5B.2 and C, n = 12 experiments, n = 24 spines, from 11 neurons and 11 mice), with only a slight but significant induction of shrinkage of the spine neck at some time points (Fig. 5C, P < 0.05, n = 19 spines, from 11 neurons and 11 mice). For comparison between the activation of one versus two clustered spines with a post-pre timing of −15 ms see Figure S8. When we analyzed the maximum uEPSP change in amplitude in each experiment and the concomitant morphological alterations in the activated spines after t-LTD induction in two clustered spines, we also observed a complete inhibition in the induction of LTD, a slight but significant reduction in spine neck length, and no changes in spine head size (Fig. 5G; uEPSP = 101.70 ± 7.02%, P = 0.81, n = 12 experiments; neck length = 85.28 ± 5.96%, P < 0.05, n = 19 spines; head volume = 102.98 ± 8.86%, P = 0.73, n = 24 spines, from 12 experiments performed in 11 neurons from 11 mice). We obtained similar results when we instead considered the average of all the values obtained following t-LTD induction in clustered spines for uEPSP amplitude, neck length and head volume (Fig. S1). These results were surprising since not only did we not observe t-LTD in clustered spines, but we also observed significant neck shrinkage with no LTP (see Fig. 1 and 3). To account for this observation, we explored if there was a correlation between the induction of plasticity in these experiments and both the shrinkage of the spine neck and the distance between the activated clustered spines – since the local voltage, and hence the induction of plasticity, could be affected by the distance between the activated clustered spines. Indeed, we found that the distance between the activated spines under these experimental conditions (t-LTD induction protocol in clustered spines) is correlated with the induction of plasticity and the shrinkage of the activated spine necks (Equation 1 in Methods; P < 0.01; Fig. S7D). This analysis suggests that during t-LTD induction the structural arrangement of clustered spines (< 40 µm) determines the sign and magnitude of the change in synaptic strength and concomitant neck shrinkage.

**Figure 5.**
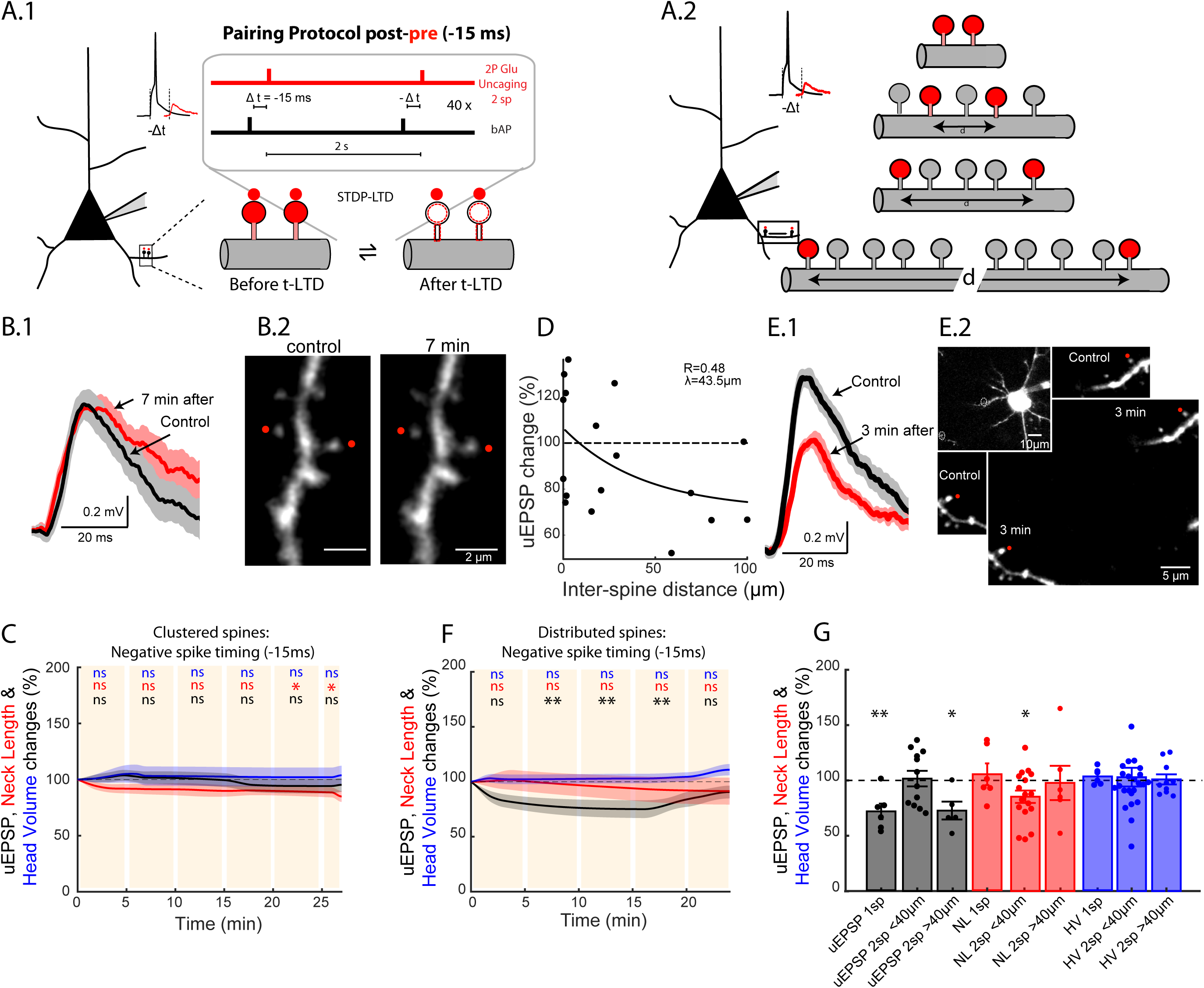
Induction of t-LTD in clustered and distributed dendritic spines: (A) Experimental protocol for the induction of t-LTD protocol (pairings of - 15 ms) in two clustered dendritic spines (< 40 µm) (A.1), or in two distributed spines (> 40 µm distance between spines). (B) An example of one experiment where two neighbouring dendritic spines were activated with a post-pre t-LTP protocol of −15 ms. (B.1) Average uEPSP recorded via a whole-cell pipette at the soma of L5 pyramidal neurons before (Control) and after the induction of t-LTD in two selected dendritic spines (red dots, B.2) located in basal dendrites. Traces in B.1 correspond to an average of 10 uEPSP generated by 2P uncaging of glutamate at the indicated spines before (black trace) and after the induction of t-LTD (red trace). (C) Time course of uEPSP amplitude (black line), the neck length (red line) and spine head volume (blue line) of the activated clustered spines after the induction of t-LTD at pairings of −15 ms. *ns*, not significant; *P < 0.05, one-way repeated measures ANOVA followed by post hoc Dunnet’s test. (D) t-LTD recovery (decrease in uEPSP amplitude, less than 100%) is dependent on inter-spine distance. Each data point represents the maximal change in uEPSP from each experiment. We use the extracted value of λ as a boundary between clustered and distributed spines. (F) Example of one experiment where two distributed dendritic spines (> 40 µm distance between spines) were activated with a post-pre timing −15 ms. (F.1) Average uEPSP response recorded via a whole-cell pipette at the soma of L5 pyramidal neurons before (Control, underneath image, F.2) and 3 min after the induction of t-LTD (F.2, over imposed image) in two selected dendritic spines located in basal dendrites. (F.2) Inset shows a low magnification image of the recorded neuron with the marked location of the selected spines. Traces in F.1 correspond to an average of 10 uEPSP generated by 2P uncaging at the indicated spines (red dots, F.2) before (black trace) and after the induction of t-LTD (red trace). (G) Time course of uEPSP amplitude (black line), the neck length (red line) and spine head volume (blue line) of the activated distributed spines (> 40 µm) after the induction of t-LTD at pairings of −15 ms. *ns*, not significant; *P < 0.05, **P < 0.01, one-way repeated measures ANOVA followed by post hoc Dunnet’s test. (H) Pooled data of the maximum changes in uEPSP amplitude (black bar and dots) and concomitant changes in neck length (red bar and dots) and head volume (blue bar and dots) of individual (1sp), clustered (2sp at < 40 µm) and distributed spines (2sp at > 40 µm) after the induction of t-LTD at a post-pre timing of −15 ms. *P < 0.05, **P < 0.01, paired t test. Time 0 in C and G is the time when the 40 pre-post repetitions of the induction protocol were completed. NL = neck length, HV = head volume.

We next investigated the mechanisms underlying the disruption of t-LTD by activating spines separated by increasingly larger distances (Fig. 5A.2). Interestingly, the induction of t-LTD in spines separated by more than 40 µm (distributed spines) was capable of recovering the generation of LTD (Fig. 5D-E). Pooled data from all experiments demonstrate that the activation of distributed spines reliably induces t-LTD (Fig. 5F, significant reduction in uEPSP amplitude, P < 0.01, n = 5 experiments from 5 neurons and 5 mice), without triggering changes in neck length or spine head size (Fig. 5E and F). When we analyzed the maximal change in uEPSP amplitude in each experiment and concomitant spine morphological changes, we saw a significant induction of t-LTD and no change in spine morphology (Fig. 5G, uEPSP = 72.86 ± 8.08%, P < 0.05, n = 5 experiments, neck length = 97.85 ± 15.47%, P = 0.89, n = 6 spines; head volume = 101.06 ± 4.59%, P = 0.82, n = 10 spines) as what was found in experiments where t-LTD was generated at pairing times of −15 ms in single dendritic spines (Fig. 2B-C). We obtained similar results when we instead considered the average of all the values obtained following t-LTD induction in distributed spines for uEPSP amplitude, neck length and head volume (Fig. S1). No significant difference was observed between the initial uEPSP amplitude for clustered versus distributed spines activated with post-pre pairings of −15 ms (EPSP: 1.06 ± 0.13 versus 1.31 ± 0.19 mV, P = 0.25; Fig. S3). For comparison between the activation of clustered versus distributed spines after post-pre pairings of −15 ms, see Figure S8.

In summary, this data shows that the induction of t-LTD at pairing times of −15 ms was disrupted when only two clustered spines (< 40 µm apart) were nearly simultaneously activated in the basal dendrites of L5 pyramidal neurons, but could be recovered if the activated spines are distributed (> 40 µm) in the dendritic tree.

### Spine calcium transients during the induction of t-LTP and t-LTD in single and clustered dendritic spines

Calcium is critical for the induction of synaptic plasticity^39, 40, 41, 42, 43^, and high or low local concentration difference in dendrites and spines are thought to be associated with gating LTP or LTD, respectively^44, 45, 46^. Therefore, to investigate the different mechanisms – with respect to local calcium accumulations – underlying the induction of t-LTP and t-LTD in single versus two clustered spines, we performed 2P calcium imaging in a region of interest (ROI) of the activated spines and their parent dendrites during STDP induction protocols throughout each of the 40 pre-post or post-pre repetitions (see Methods). The “before” images correspond to the calcium signals observed in the ROI right before the pairing in each repetition – uncovering the lack or presence of local calcium accumulation during the 40 pairing repetitions. The “after” images correspond to the calcium signals observed in the ROI right after the pairing in each repetition – uncovering a proxy for the amplitude and local calcium accumulation during the 40 pairing repetitions.

To dissect potential differences in local calcium signals and accumulation that can account for the presence or absence of t-LTP and t-LTD induction in clustered versus distributed spines, we imaged 2P calcium activity during five different STDP induction protocols: (1) pre-post pairing of +7 ms in one spine; (2) pre-post pairing of +7 ms in two clustered spines; (3) pre-post pairing of +13ms in one spine; (4) post-pre pairing of −15 ms in one spine; (5) post-pre pairing of −15 ms in two clustered spines.

During the pre-post (+7 ms) pairing protocol in single spines we found that, across the 40 repetitions, there was little to no calcium accumulation in the spine or dendrite (Fig. 6A and left panels in Fig. 6B, n = 7 experiments, from 6 neurons, and 4 mice). As expected, there was, however, a significant increase in calcium immediately following the stimulation (left panels in Fig. 6D) that due to the lack of accumulation throughout the 40 repetitions, did not build up a local calcium signal in the activated dendrites and spines. In contrast, when we applied the exact same pairing protocol (pre-post + 7ms) in two clustered spines, there was a striking calcium accumulation in both the activated spines and dendrite that was evident when we analyzed the images taken before (Fig. 6C and middle panels in Fig. 6B) and after stimulation (middle panels in Fig. 6D, n = 6 experiments, from 4 neurons, and 4 mice). Thus, activating just one additional spine using the same pairing protocol alters the calcium dynamics (compare black and green traces in right panels of Fig. 6B and D), possibly through a mechanism that is incapable of extruding calcium increases in spines in between pre-post repetitions, leading to its build up in spines and parent dendrites, which ultimately guide the induction of plasticity.

**Figure 6.**
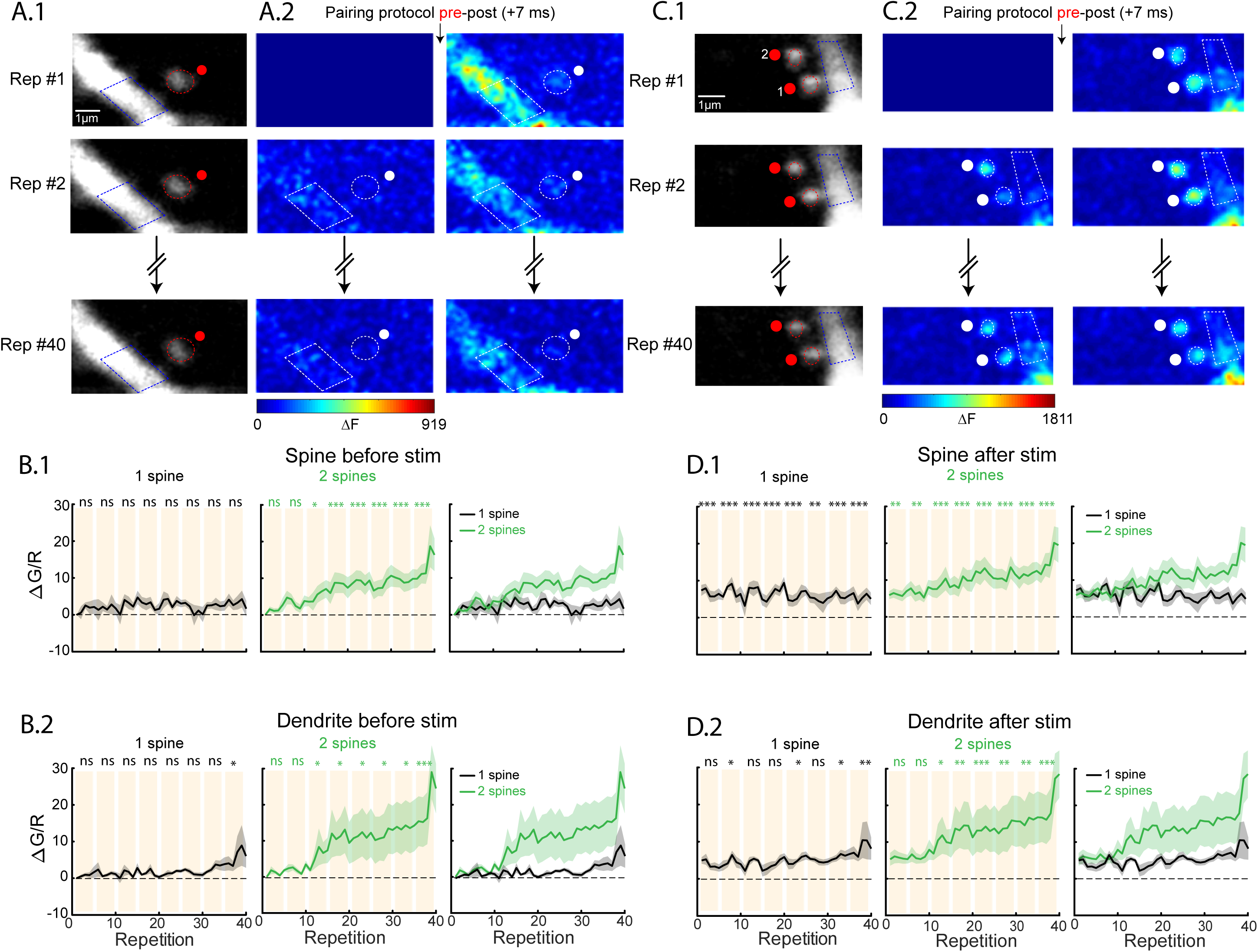
Calcium dynamics in single and two clustered spines during a pre-post pairing protocol of + 7ms. (A.1) Single 2P images of a spine and dendrite from a L5 pyramidal neuron loaded with Alexa 594 (100uM) and Fluo4 (300uM). Red ellipses and blue polygons indicate the ROIs selected for the calcium signal analysis. (A.2) Two photon calcium signal images before (left panels) and after (right panels) a pre-post pairing protocol (+7 ms). The 1^st^, 2^nd^, and 40^th^ repetitions of the pairing protocol are shown here. The change in calcium fluorescence from baseline (ΔF) is color coded. Only positive changes in fluorescence are shown. (B) Population averages of the calcium signals 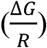 measured in spines (B.1) and dendrites (B.2) before the pairing protocol performed in 1 spine (left panels) and 2 spines (middle panels). The right panels shows the superimposed 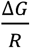 population averages in 1 spine (black lines) and 2 spines (green lines). (C.1) Single 2P images of two clustered spines and dendrite from a L5 pyramidal neuron loaded with Alexa 594 (100µM) and Fluo4 (300µM). Red ellipses and blue polygons indicate the ROIs selected for the calcium signal analysis. (C.2) Two photon calcium signal images before (left panel) and after (right panels) a pre-post pairing protocol (+ 7ms). The 1^st^, 2^nd^, and 40^th^ repetitions of the pairing protocol are shown here. The change in calcium fluorescence from baseline (ΔF) is color coded. Only positive changes in fluorescence are shown. (D) Population averages of the calcium signals 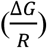 measured in spines (D.1) and dendrites (D.2) after the pairing protocol performed in one spine (left panels) and two spines (middle panels). The right panels shows the superimposed 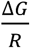 population averages in one spine (black lines) and two spines (green lines). *ns*, not significant; *P < 0.05; **P < 0.01; ***P < 0.001, one-way repeated measures ANOVA followed by post hoc Dunnet’s test.

Since the activation of a single spine with a repetitive pre-post pairing protocol of +13 ms reliably induced t-LTP (Fig. 1B-C), we performed experiments to measure the calcium signals in the activated spine and parent dendrite during the pre-post (+13 ms) pairing protocol. We found that, across the 40 repetitions, there was a significant calcium accumulation in both the activated spine and dendrite that was evident when we analyzed the images taken before (Fig. 7A and B) and after stimulation (Fig. 7A and C).

**Figure 7.**
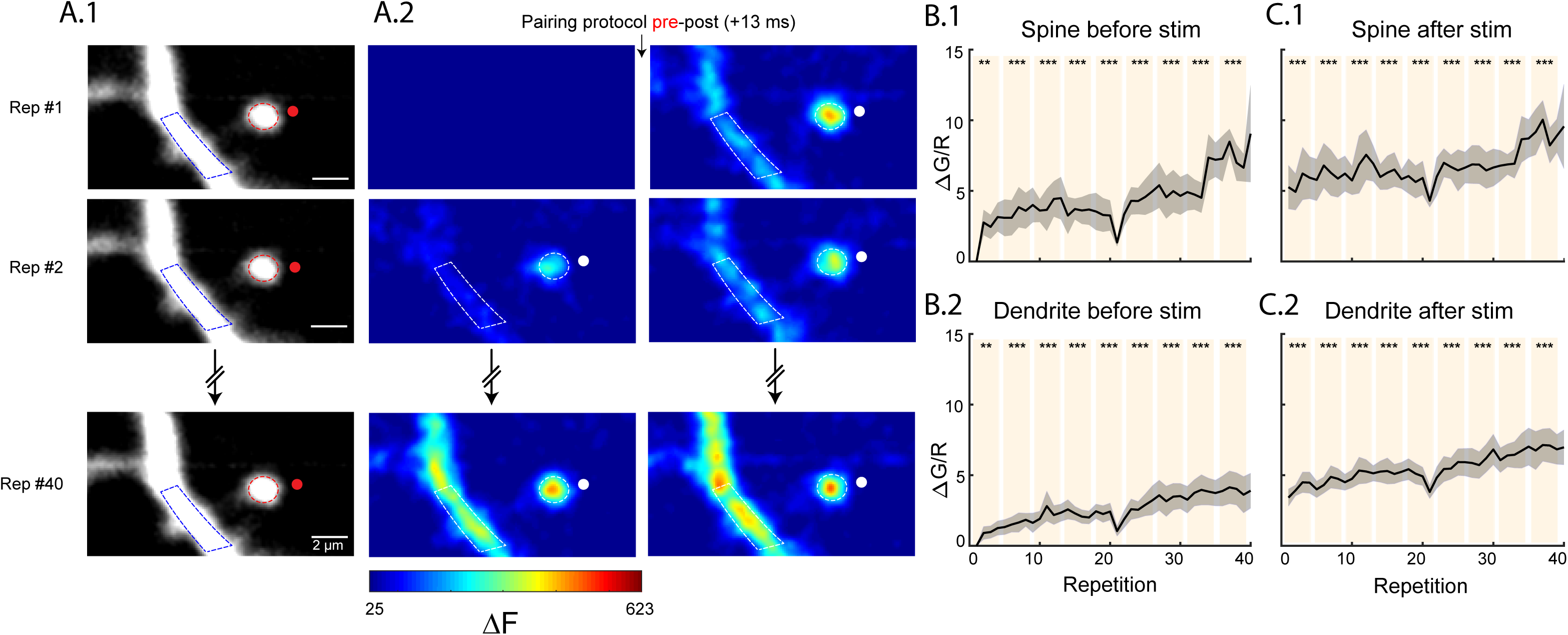
Calcium dynamics in single spines during a pre-post pairing protocol of +13ms. (A.1) Single 2P images of a spine and dendrite from a L5 pyramidal neuron loaded with Alexa 594 (100µM) and Fluo4 (300µM). Red ellipses and blue polygons indicate the ROIs selected for the calcium signal analysis. (A.2) Two photon calcium signal images before (left panels) and after (right panels) a pre-post pairing protocol of +13 ms. The 1^st^, 2^nd^, and 40^th^ repetitions of the pairing protocol are shown here. The change in calcium fluorescence from baseline (ΔF) is color coded. Only positive changes in fluorescence are shown. (B) Population averages of the calcium signals 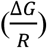 measured in spines (B.1 and C.1) and dendrites (B.2 and C.2) before (B) and after the pairing protocol. *ns*, not significant; *P < 0.05; **P < 0.01; ***P < 0.001, one-way repeated measures ANOVA followed by post hoc Dunnet’s test.

In summary, the induction of t-LTP in a single spine at a pre-post pairing protocol of +13 ms (Fig. 7) and in clustered spines at a pairing time of +7ms (which is otherwise ineffective in triggering t-LTP if only one spine is being activated) (Fig. 6) are correlated with an accumulation of calcium in spines and parent dendrites, likely being responsible for the induction of t-LTP. Next, we performed the same experiments with a post-pre (−15 ms) pairing protocol in both single and clustered spines. In single spines, we observed moderate calcium increases (Fig. 8A, n = 5 experiments, from 4 neurons, and 4 mice) that were observed when we analyzed images taken before (left panels in Fig. 8B) and after the post-pre stimulation (left panels in Fig. 8D).

**Figure 8.**
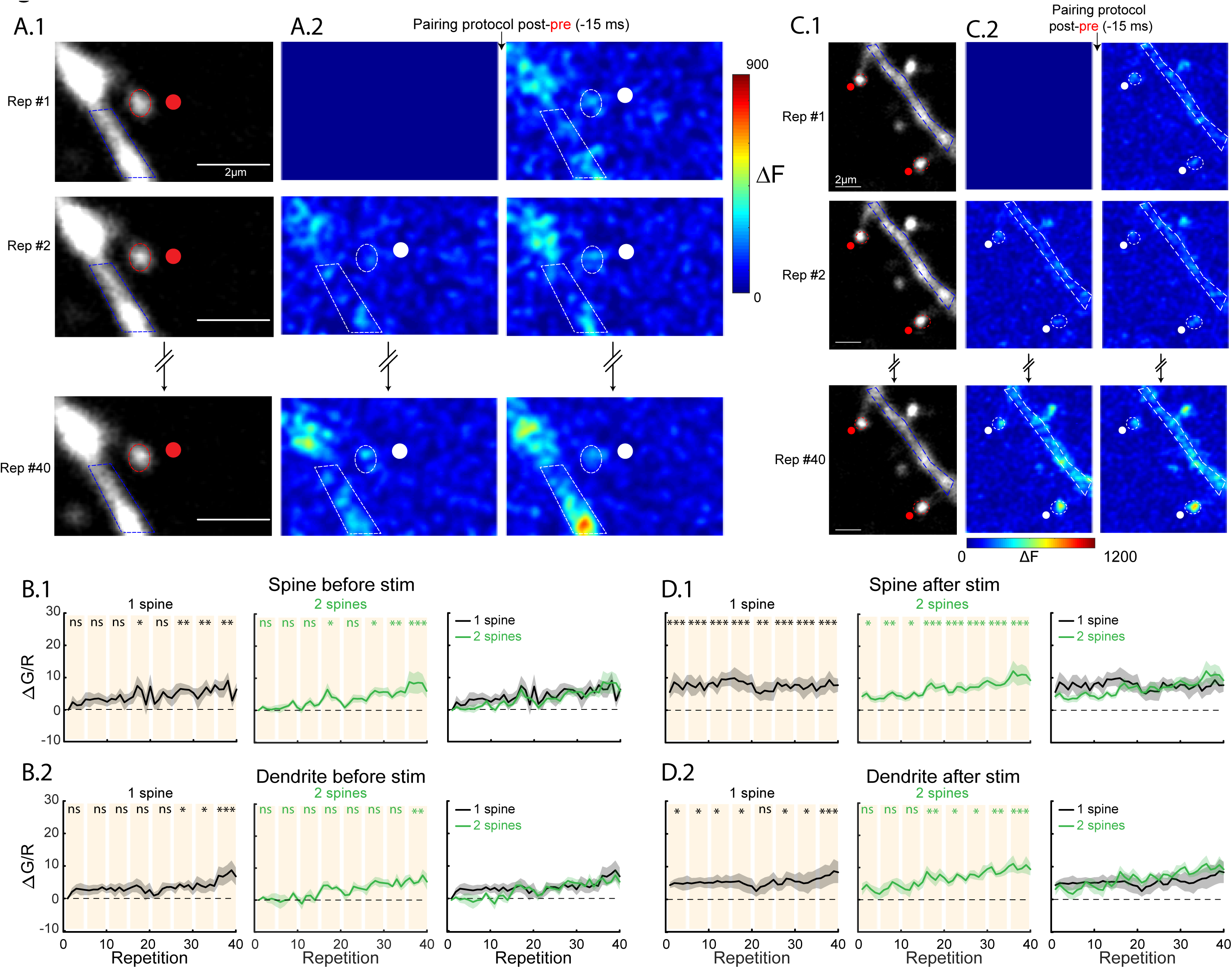
Calcium dynamics in single and two clustered spines during post-pre pairing protocol. (A1) Single 2P images of a spine and dendrite from a L5 pyramidal neuron loaded with Alexa 594 (100µM) and Fluo4 (300µM). Red ellipses and blue polygons indicate the ROIs selected for the calcium signal analysis. (A.2) Two photon calcium signal images before (left panels) and after (right panels) a post-pre pairing protocol (−15 ms). The 1^st^, 2^nd^, and 40^th^ repetitions of the pairing protocol are shown here. The change in calcium fluorescence from baseline (ΔF) is color coded. Only positive changes in fluorescence are shown. (B) Population averages of the calcium signals 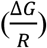 measured in spines (B.1) and dendrites (B.2) before the pairing protocol performed in one spine (left panels) and two spines (middle panels). The right panels shows the superimposed 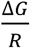 population averages in one spine (black lines) and two clustered spines (green lines). (C.1) Single 2P images of two clustered spines and dendrite from a L5 pyramidal neuron loaded with Alexa 594 (100µM) and Fluo4 (300µM). Red ellipses and blue polygons indicate the ROIs selected for the calcium signal analysis. (C.2) Two photon calcium signal images before (left panel) and after (right panels) a pre-post pairing protocol (−15 ms). The 1^st^, 2^nd^, and 40^th^ repetitions of the pairing protocol are shown here. The change in calcium fluorescence from baseline (ΔF) is color coded. Only positive changes in fluorescence are shown. (D) Population averages of the calcium signals 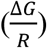 measured in spines (D.1) and dendrites (D.2) after the pairing protocol performed in one spine (left panels) and two spines (middle panels). The right panels shows the superimposed 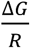 population averages in 1 spine (black lines) and 2 spines (green lines). *ns*, not significant; *P < 0.05; **P < 0.01; ***P < 0.001, one-way repeated measures ANOVA followed by post hoc Dunnet’s test.

Surprisingly, we found similar results to those observed with single spine t-LTD induction protocols, when we applied the same pairing protocol in two clustered spines (Fig. 8C, n = 6 experiments, from 4 neurons, and 3 mice) before (middle panels in Fig. 8B) or after the induction protocol (middle panels in Fig. 8D), even though no plasticity is induced in this condition. As suggested by previous studies^47^, we hypothesize that the range of spine calcium levels required for the induction of t-LTD is relatively narrow, and that the resolution with our current experimental set-up is not sufficient to tease apart significantly different calcium dynamics in one versus two clustered spines during a post-pre pairing protocol of −15 ms. Moreover, models of STDP provide evidence that, in addition to overall calcium levels, the detailed time course of calcium levels in the postsynaptic cell during a pairing protocol also guide the induction of plasticity^48^. Nonetheless, these results suggest that the induction of t-LTD does not require significant calcium accumulations during the 40 repetitions, and most likely depends on the amplitude of calcium signals right after the stimulation.

## DISCUSSION

We uncovered the STDP rules for single, clustered and distributed dendritic spines in the basal dendrites of L5 pyramidal neurons. Our results show that the induction of STDP in single spines follows a classical Hebbian STDP learning rule that is bidirectional, in which presynaptic input leading postsynaptic spikes generates t-LTP and postsynaptic spikes preceding presynaptic activation of single dendritic spines results in t-LTD. Furthermore, we found that the induction of t-LTP triggers the shrinkage of the activated spine neck, without any significant changes in the spine head size, building upon our previous findings of the activity-dependent shrinkage of the spine neck^2^. Our results indicate that the induction of t-LTP requires 1) the incorporation of new GluR-1 receptors with PDZ-domain containing proteins in the PSD and, 2) an actin polymerization-dependent neck shrinkage of the activated spine neck (Fig. 4). We showed that the induction of t-LTP triggers actin-dependent neck shrinkage, which is likely required for the lateral diffusion of GluR-1 receptors from the spine neck to the spine head, and its incorporation to the PSD – generating plasticity. In support of this spine mechanism of LTP induction is a recent report showing that AMPA receptor surface diffusion is fundamental for the induction of hippocampal LTP and contextual learning^49^. In addition, we found that the induction of t-LTD was not accompanied with spine neck or head changes, which is at odds with previous findings suggesting structural changes in spine head volume during the induction of LTP or LTD^4, 8, 50^. The discrepancy between our results and those observed previously after the induction of t-LTP (head enlargement^7, 50^), LTP^4^, or LTD (head shrinkage,^8^) using glutamate uncaging are likely explained by methodological differences. While our data was obtained using ACSF with physiological concentrations of magnesium and calcium, those from other reports were done in low or a magnesium-free ACSF^4, 8^, low calcium extracellular solution for the induction of LTD^8^, or in a magnesium-free ACSF and an intracellular solution containing 5 µM actin that was required for the t-LTP-mediated spine head enlargements^50^.

Nonetheless, it has been shown in vivo that a spike-timing protocol triggers receptive field plasticity in layer 2/3 pyramidal neurons is correlated with spine head volume changes (enlargement and shrinkage) observed after 1.5-2 hours^51^. Taken together, these data, and previous observations^2^, suggest that there is a new form of structural spine plasticity during t-LTP that involves rapid neck shrinkage without head volume enlargements. In addition, we show that the induction of t-LTD does not require structural spine changes. Although spines have the machinery and do undergo structural head changes, we propose that our results represent a stage during memory formation that occurs before structural head volume changes, a process likely linked with memory consolidation. Importantly, our data suggest that during STDP, the use of spine volume changes as the sole proxy for LTP or LTD^51^ is not a complete representation of plasticity in spines from dendrites in cortical pyramidal neurons.

We then explored the functional consequences of synaptic cooperativity of nearly simultaneous excitatory inputs on STDP. Our results show that the induction of t-LTP in two clustered spines - separated by less than 5 µm - is sufficient to induce LTP and shrinkage of the activated spine necks at a pre-post timing (+ 7ms) that is otherwise ineffective at triggering significant morphological changes and synaptic potentiation when only one spine is being activated. To uncover the distance between the nearly simultaneously activated spines capable of supporting synaptic cooperativity and the induction of t-LTP, we varied the distance of the activated spines. Our results show that the induction of t-LTP is suppressed when spines are separated by more than 5 µm apart (Fig. 3 and Fig. S5A), with an effective length constant (λ), that represents the length at which the electrotonic potential decays to a value of 37% of that at the point of origin, of 5.7 µm These results show that the nearly simultaneous activation of clustered spines, less than < 5 µm apart, can extends the pre-post timing window that can trigger potentiation. This data, together with results showing that the induction of t-LTP in one spine can change the threshold for the induction of plasticity in a spine located at less than 10 µm away and activated 90 seconds later^7^, indicate that there is a spatially restricted dendritic compartment that is required for spine crosstalk and the broadening of the STDP timing required for triggering t-LTP (Fig. 9).

**Figure 9.**
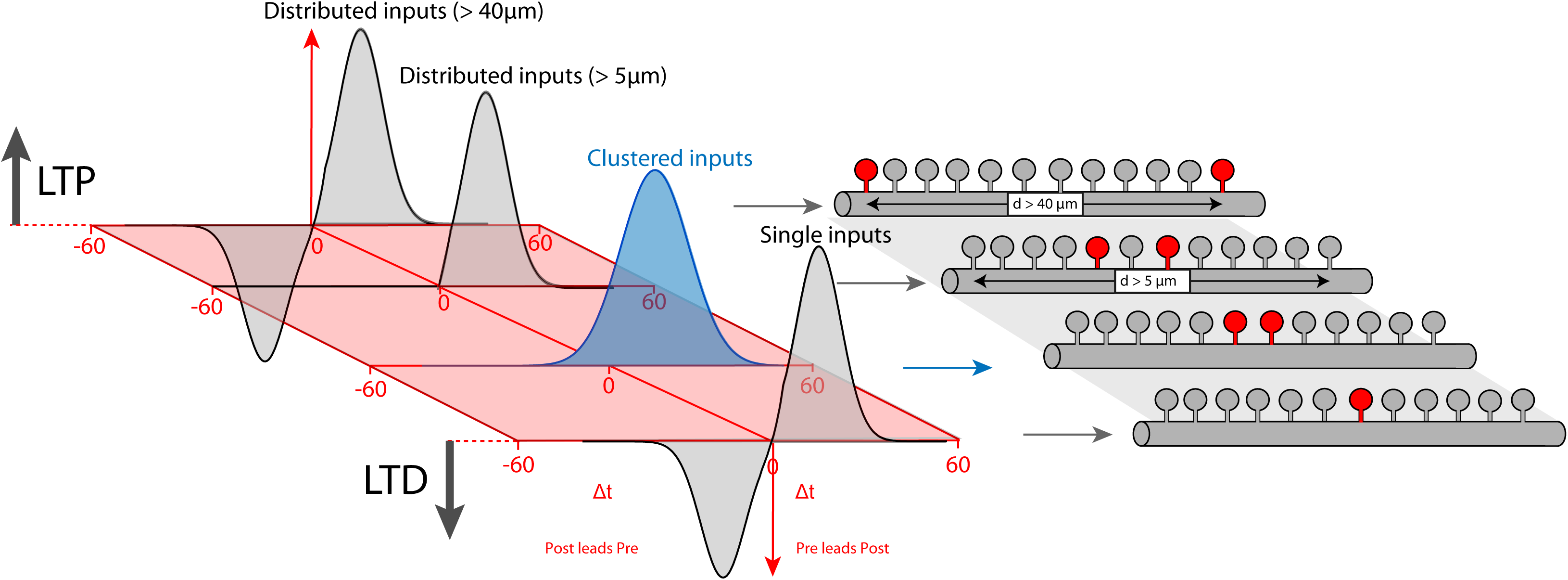
STDP learning rule for single distributed and clustered dendritic spines. STDP learning rule in the basal dendrites of L5 pyramidal neurons as a function of the structural organization of excitatory inputs in basal dendrites of L5 pyramidal neurons. Note how STDP in single, or distributed spines (separated by > 40 µm), follow a bidirectional Hebbian STDP rule. Importantly, our model indicate that the induction of t-LTD can be disrupted by the co-activation of two clustered spines separated by < 40 µm, and t-LTP enhanced by the co-activation of two clustered spines separated by < 40 µm by synaptic cooperativity. We propose that synaptic cooperativity generates local dendritic depolarization high enough to disrupt bidirectional STDP, leading to STDP that only encompasses LTP.

On the other hand, the induction of t-LTD in two clustered spines disrupts the generation of LTD leading to a STDP learning rule that is incapable of supporting LTD, but only encompasses LTP (Fig. 9). We next investigated the dendritic mechanisms responsible for the disruption of t-LTD, and found that the induction of t-LTD is recovered when the activated spines are separated by more than 40 µm (Fig. 5, Fig. 9 and Fig. S5B). Interestingly, the effective length constant (λ) in the basal dendrites of L5 pyramidal neurons has been reported to be 50 µm^52^. This value of λ supports the idea that significant voltage attenuations – capable of recovering LTD – can be expected when the t-LTD induction protocol is triggered in spines that are separated by more than 40 µm in the basal dendrites of L5 pyramidal neurons (Fig. 5D-G). However, we cannot discard that other mechanisms, such as the diffusion of active molecules^5^, could contribute to the switch from t-LTD to no-LTD induction observed in distributed/single spines and clustered spines, respectively. These results are in discrepancy with observations showing that in connected pairs of L5-L5 pyramidal cells, t-LTD is reliably generated after post-pre pairing protocols^13^. A likely explanation for this apparent controversy is that the synaptic inputs from one L5 pyramidal neuron to another are distributed^53^. Importantly, clustered and distributed excitatory inputs have been described in the dendrites of pyramidal neurons both *in vitro* and *in vivo*^1, 54–56^. Our results clearly show the functional importance that the structural and temporal organization of excitatory synaptic inputs have on the induction of t-LTP and t-LTD, and how just two clustered excitatory synaptic inputs are capable of altering the STDP learning rule in the basal dendrites of L5 pyramidal neurons (Fig. 9).

How the synaptic activation of just one extra clustered spine is capable of (1) inducing t-LTP at a pre-post timing that is otherwise ineffective in inducing potentiation and (2) disrupting the induction of t-LTD? To explore the mechanisms that may be responsible for these observations we imaged local calcium signals in the activated spines and parent dendrites before and after each of the 40 pairings performed during t-LTP and t-LTD induction protocols. Our reasoning for performing these experiments was based on findings that different levels of depolarization gate local calcium signals, which depending on its magnitude and kinetics, can generate LTP (high calcium) or LTD induction (sustained but moderate calcium signals)^10, 44, 57^. In addition, calcium-based modeling studies of STDP have shown that different calcium dynamics mediate the induction of t-LTP versus t-LTD^47, 48^. Specifically, the calcium control hypothesis indicates that large levels of calcium (above a plasticity threshold, ϴ_p_) are thought to lead to t-LTP whereas more moderate, prolonged levels (between the depression threshold, ϴ_dSTART_, and ϴ_dSTOP_)) give rise to t-LTD and a mid-level range in which t-LTD does not occur (below ϴ_dSTART_)^48, 58, 59^. A major assumption of these models is infinite time constants for synaptic variables at resting calcium levels so that the synaptic changes do not to decay after the presentation of the stimulus^47^ - a significant constraint for the stabilization of synaptic changes. A potential solution to this problem is the degree of local calcium accumulation observed in the activated spines throughout the t-LTP or t-LTD induction protocol. In fact, these models are consistent, fundamentally, with our results which show that a pre-post pairing of +7 ms protocol in two clustered spines, and a pre-post pairing of +13 ms protocol in a single spine gives rise to t-LTP accompanied by a substantial increase in the intracellular calcium levels following each pairing repetition, and a significant accumulation of local calcium levels throughout the induction protocol – likely mediated by the inability to efficiently extrude the local calcium signals in between each pre-post pairing at this induction frequency (Fig. 6 and 7). We propose that the local spine calcium accumulation we observe provides a new and key variable for the induction of plasticity, which reduces the constraints imposed by calcium-base models for the stabilization of synaptic changes^47, 48, 58, 59^.

These changes in local spine and dendritic calcium signals observed when clustered spines are activated at pairings of + 7ms (Fig. 6) or in a single spine activated at pairings of + 13ms (Fig. 7) suggest that perhaps ϴ_p_ can be reached only with ∼ 10-20 pre-post pairings (∼20-40 seconds). In contrast a pairing protocol of + 7ms in one spine induces no plasticity, and calcium levels that are effectively extruded in between pairings leading to no calcium accumulation during the induction protocol (reaching levels below ϴ_p_ and ϴ_dSTART_). These results suggest that it is not only the amplitude of the local calcium signals after each pairing, but also the local calcium accumulation during the induction protocol (40 pairings, ∼80 seconds) in spines and dendrites that are required to reach ϴ_p_ for the induction of t-LTP in clustered spines. As mentioned before, recently it has been demonstrated in vivo that spike timing-induced receptive field plasticity, with millisecond time delays between visual stimulus (pre) and optogenetic stimulation in layer 2/3 pyramidal neurons (post), is correlated with increases in synaptic strength^51^. These results together with evidence from other in vivo studies showing that layer 5 pyramidal neurons can spike up to frequencies of 20 Hz during movement^60^, suggest that our pairing protocol, and findings, are likely present under certain in vivo conditions and are relevant for plasticity of networks and ultimately behaviour.

Our results further show that a post-pre protocol of −15 ms in a single spine induces t-LTD and moderate intracellular calcium signals in spines and parent dendrites after each pairing, without an evident increase in local calcium accumulation. These results possibly reflect that the calcium signal generated during the induction protocol passed ϴ_dSTART_ and remained for several seconds in this permissive calcium concentration window – between ϴ_dSTART_ and ϴ_dSTOP_ – generating LTD. Activating two clustered spines with the same protocol, however, does not induce plasticity and gives rise to an apparent smaller initial calcium accumulation than that observed with the activation of a single spine but with a slow build-up of calcium. These results possibly reflect that the spine calcium levels crossed ϴ_dSTART_ only after > 20 repetitions and then crossed ϴ_dSTOP_ and ϴ_pSTART_ after a few (<10) repetitions reaching slightly higher local calcium levels. Further experiments to prove this calcium control hypothesis of t-LTD induction, with better temporal detection tools are needed.

These findings presented here are quite remarkable since stimulating just one additional spine during a STDP protocol can alter the calcium dynamics and the induction of t-LTP and t-LTD. To our knowledge, this is the first demonstration of the functional relevance that the structural organization and nearly simultaneous subthreshold activation of only a few clustered inputs in the dendrites of pyramidal neurons have on plasticity. We propose the term *micro clusters* to describe this structural and functional modality of synaptic connectivity. In fact, the relevance of synaptic *micro clusters* on the input/output properties of pyramidal neurons is also supported by three dimensional electron microscopy and neuronal reconstruction studies that have shown the presence of postsynaptic innervation of the same axon spaced at less than 10 µm in the basal dendrites of L2/3 pyramidal neurons from the medial entorhinal cortex^54^, L5 pyramidal neurons from somatosensory cortex^55^ and in the distal apical tuft dendrites in stratum lacunosum-moleculare of hippocampal CA1 pyramidal neurons^56^. In addition to having spines innervated by the same axon, it is likely that functional synaptic *micro clusters* can be gated by the convergence of different axons, which could increase the computational power of cortical circuits through a multi-neuronal control of synaptic cooperativity and ultimately the implemented STDP learning rule. Furthermore, it has been shown that orientation selectivity in visual cortex is correlated with the degree of spatial synaptic clustering of co-tuned synaptic inputs within the dendritic field^61^, and that functional clusters of dendritic spines separated by less than 10 µm share similar spatial receptive field properties, spontaneous and sensory-driven activity^62^. Interestingly, is has been recently shown in the mouse visual cortex that a single axon can contact clustered synapses in the postsynaptic neuron, but that local clustering is not favoured over widespread spacing of synaptic inputs^63^. In addition, recently it has been demonstrated in L2/3 pyramidal neurons from mouse primary visual cortex that synaptic *micro clusters* reflects the interaction of functionally similar inputs - with similar orientation preference - from different sources – callosal and non-callosal inputs^64^, providing functional and anatomical data for the presence of multi-neuronal control of synaptic cooperativity in a short dendritic segment. Moreover, it has been shown that coactive neighboring synapses drive the maturation of cluster synapses in CA1 pyramidal neurons^65^, suggesting that *micro clusters* also play an important role in the development and shaping of network connectivity. Taken together these reported findings and our data suggest that the functional specificity and structural arrangement of synaptic inputs, distributed or forming *micro clusters* in the dendrites of pyramidal neurons, are fundamental for guiding the rules for sensory perception, affecting the STDP learning rule, learning and memory, and ultimately cognition.

## METHODS

### Brain slice preparation and electrophysiology

Brains from postnatal day 14-21 C57B/6 mice - anesthetized with isoflurane - were removed and immersed in cold (4°C) oxygenated sucrose cutting solution containing (in mM) 27 NaHCO_3_, 1.5 NaH_2_PO_4_, 222 Sucrose, 2.6 KCl, 1 CaCl_2_, and 3 MgSO_4_. Coronal brain slices (300-µm-thick) of visual cortex were prepared as described^23^. Brain slices were incubated for 1/2 hour at 32°C in artificial cerebrospinal fluid (ACSF, in mM: 126 NaCl, 26 NaHCO_3_, 10 Dextrose, 1.15 NaH_2_PO_4_, 3 KCl, 2 CaCl_2_, 2MgSO_4_) and then transferred to a recording chamber. Electrophysiological recordings were performed in whole-cell current-clamp configuration with MultiClamp 700B amplifiers (Molecular Devices) in L5 pyramidal neurons with a patch electrode (4-7 MΩ) filled with internal solution containing (in mM) 0.1 Alexa Fluo 568, 130 Potassium D-Gluconic Acid (Potassium Gluconate), 2 MgCl_2_, 5 KCl, 10 HEPES, 2 MgATP, 0.3 NaGTP, pH 7.4, and 0.4% Biocytin. All experiments were conducted at room temperature (∼20-22°C). We did not extend our experiments to include voltage-clamp recordings since recent evidence indicates that the high spine neck resistance can prevent the voltage-clamp control of excitatory synapses and that these measurements can be significantly distorted in spiny neurons^66^.

### Two-photon imaging and two-photon uncaging of glutamate

Two-photon imaging was performed with a custom-built two-photon laser scanning microscope, consisting of 1) a Prairie scan head (Bruker) mounted on an Olympus BX51WI microscope with a 60X, 0.9 N.A. water immersion objective; 2) a tunable Ti-Sapphire laser (Chameleon Ultra-II, Coherent, >3 W, 140-fs pulses, 80 MHz repetition rate), 3) two photomultiplier tubes (PMT) for fluorescence detection. Fluorescence images were detected with Prairie software (Bruker).

Fifteen minutes after break-in, two-photon scanning images of basal dendrites were obtained with 720 nm and low power (∼5 mW on sample (i.e., after the objective)) excitation light and collected with a PMT. Two-photon uncaging of 4-methoxy-7-nitroindolinyl (MNI)-caged L-glutamate (2.5 mM; Tocris) was performed as previously described^67^. This concentration of MNI-glutamate completely blocked IPSCs^68^, thus, our results represent excitatory inputs only. Uncaging was performed at 720 nm (∼25-30 mW on sample). Note that the laser power used for imaging is not sufficient to result in any partial uncaging of glutamate (Fig. S9). Activated spines were mostly on the second and third branch of the basal dendrites and were on average 40.22 ± 1.62 µm away from the soma (range of distances: ∼10 to 80 µm from soma, n = 104 spines, Fig. S10). Only spines with a clear head contour and that were separated by >1 µm from neighboring spines were selected. The location of the uncaging spot was positioned at ∼ 0.3 µm away from the upper edge of the selected spine head (red dot in figures), which has a spatial resolution of 0.71 and 0.88 µm for one and two spines respectively (Fig. S11). Care was taken to maintain the position of the uncaging spot. After each uncaging sequence, the spot was repositioned to keep the same distance of ∼ 0.3 µm from the edge of the soma and to avoid artificial potentiation or depression. The uncaging-induced excitatory postsynaptic potentials (uEPSP) were recorded at the soma of L5 pyramidal neurons. Importantly, the kinetics of uEPSPs from the present study are not significantly different (10/90 rate of rise: 0.07 ± 0.014 mV/ms, p=0.92; duration: 115.5 ± 15.3 ms, p=0.65) from those triggered at 37°C^23^.

### Spike-timing-dependent plasticity (STDP) induction protocol

To induce t-LTP in single spines, we used two-photon uncaging of MNI-glutamate (40 times every 2 seconds, with each uncaging pulse lasting 2 ms), which, after 7 or 13 ms, was followed by a backpropagating action potential (bAP) (triggered by a brief (10 ms) current injection (0.4-0.6 nA) in the soma). To induce t-LTD in single spines, two-photon uncaging of MNI-glutamate (40 times every 2 seconds, with each uncaging pulse lasting 2 ms) was preceded for −15 or −23 ms by bAP. When we evaluated t-LTP and t-LTD in two spines, we used similar protocols to those described above, but the spines were activated with two-photon uncaging of MNI-glutamate sequentially with an onset delay of ∼2.1 ms for the second spine. No significant difference was observed in the in 10/90 rise time of the uEPSPs triggered when one versus two spines were activated (9.05 ± 1.19 ms versus 9.49 ± 0.54 ms, respectively; p = 0.71).

To evaluate the morphological and synaptic strength of the activated spines before and after the STDP induction protocol, we performed 2P imaging, and low frequency 2P uncaging (0.5 Hz) in single or multiple spines. To establish the time course of the changes in uEPSP amplitude, neck length and head volume following STDP induction, for each experiment, we interpolated the data taken at different time points using the *interp1* function in MATLAB (MathWorks) with the *pchip* option, which performs a shape-preserving piecewise cubic interpolation. Note that we constrained this fit so that it terminated with a slope of zero following the last data point. Then, for each condition, we averaged the uEPSP amplitude, neck length and head volume temporal traces. The length of the x-axis was set as the time at which the last data point was obtained for those sets of experiments. Shaded area represents ±SE. To determine at which time the uEPSP amplitude, neck length and head volume temporal traces are significantly different from baseline, we binned the temporal traces every 5min and tested whether it was significantly different from baseline (100%).

The time at which the maximal change in uEPSP was observed after t-LTP and t-LTD induction was used to calculate the percent change from control, and the percent changes in neck length and head volume. These analyses are displayed in Figures 1 to 5.

### Experimental checkpoints and data analysis

Electrophysiological data were analyzed with Wavemetrics software (Igor Pro) and MATLAB. The resting membrane potential of the recorded L5 pyramidal neurons was −60.25 ± 0.54 mV (recorded from a random sample of n = 58 neurons from a total of n= 99 neurons tested), which is similar to what others have reported in acute mouse brain slices of a similar age^69^. After taking this measurement, pyramidal neurons were maintained at – 65 mV in current clamp configuration throughout the recording session. Only neurons for which the injected current to hold the cell at – 65 mV was < 100 pA were included in this study. For the generation of bAP, only action potentials (AP) with amplitude of > 45 mV from threshold to the peak amplitude were considered for analysis. We found a mean AP amplitude from threshold of 63.14 ± 1.73 mV before STDP induction which was similar to that measured during the STDP protocol (63.17 ± 1.76 mV) and at all time points after the STDP protocol (62.08 ± 1.91, pooled data of AP amplitude measured at all time points from each experiment, p = 0.93; n = 58). No significant difference was found by comparing AP amplitude before and after the STPD protocol (p > 0.5, paired t-test, n=58). The absolute AP amplitude, measured from the membrane potential to the peak depolarization amplitude, before the STDP protocol was 79.47 ± 2.21 mV (n = 58), and remained stable after the STDP protocol (80.11 ± 1.4 mV, pooled from all times after STDP, p = 0.93).

In most experiments, two control tests (each consisting of 10 uncaging pulses at 0.5 Hz), spaced by 5 min were performed to assess the reliability of the uEPSP amplitude. Only experiments for which uEPSP amplitudes were not significantly different before and after 5 minutes in control conditions were considered for analysis (less than 10% variation).

Synaptic plasticity was assessed by two parameters: the uEPSP amplitude and the spine morphology (neck length and head volume). The peak uEPSP amplitude was measured from each individual uEPSP by taking the peak value and averaging 2 ms before and after using Wavemetrics (Igor Pro). Only uEPSPs that were > 0.1 mV when one spine was activated in the control condition (i.e., before the induction of plasticity) were included in the analysis.

Synaptic plasticity was determined by the relative change of uEPSP amplitude (average of 10 uEPSP) measured before and after the STDP protocol. For each experiment, we evaluated whether the STDP protocol generated potentiation or depression by determining how many uEPSP data points fell above or below baseline values over the course of the experiment. Potentiation was defined as the majority of uEPSP amplitude data points measured over time increasing relative to baseline, and the maximum uEPSP increase was used for statistical test. Similar results were obtained when we instead considered the average uEPSP change following STDP induction for each experiment. Depression was defined as the majority of uEPSP amplitude data points measured over time decreasing relative to baseline, and the maximum uEPSP decrease was used for statistical test. Similar results were obtained when we instead considered the average uEPSP change following STDP induction for each experiment. The spike timings +7 ms and +13 ms (pre leads post) or −23 ms and −15 ms (post leads pre) correspond to the delta time offset between the beginning of the uncaging pulse (pre) and the beginning of the bAP pulse (post) repeated 40 times.

The analysis of spine morphology was performed from z-projections of the whole spine using ImageJ^70^ (neck length) and MATLAB (MathWorks) (head volume). The neck length was measured from the bottom edge of the spine head to the edge of its parent dendritic shaft using the segmented line tool in ImageJ. We selected mostly spines with a spine neck longer than 0.2 µm. For those with a shorter neck, we did not report their length for analysis and statistics due to the diffraction limited resolution of our images. For spines whose necks shrunk after the STDP protocol below the diffraction limited resolution of our microscope, we set their length as the minimal measurement of spine neck length reported by Tonnesen et al., using STED microscopy (0.157 µm)^17^. We estimated the relative spine head volume using the ratio of the maximum spine fluorescence and the maximum fluorescence observed in the dendrite measured from z-projections of the whole spine^71, 72^. To obtain the spine volume, we then multiplied this ratio by the PSF of our microscope (0.11 fL)^73^. Linear optimization techniques were used to determine the correlation between EPSP change, neck length change and distance between 2 activated spines. Specifically, the change in uEPSP amplitude was modeled using the following equation:

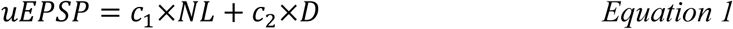

Where *uEPSP* and *NL* are the percent change in uEPSP and neck length, respectively, following the STDP protocol, *D* is the distance between the 2 spines, and *c_1_*, *c_2_*and *c_3_* are constant coefficients. These parameters were estimated using a least squares technique to obtain an optimal fit of the data that minimized the sum of the residuals squared. The relationship between inter-spine distance and the percent change in uEPSP was fit with the following exponential equation:

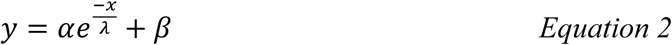

where α, β and λ are constants, y represents the change in uEPSP, and x is the inter-spine distance.

### Calcium imaging

During calcium imaging experiments, we performed whole-cell current-clamp recordings of L5 pyramidal neurons with a patch electrode containing calcium indicator Fluo-4 (300 µM; Thermo Fisher) and Alexa-594 (100 µM) diluted in an internal solution containing (in mM) 130 D-Gluconic Acid, 2 MgCl_2_, 5 KCl, 10 HEPES, 2 MgATP, 0.3 NaGTP, pH 7.4, and 0.4% Biocytin. To perform sequential 2P calcium imaging and 2P uncaging of caged glutamate in selected spines at one wavelength (810 nm), we used ruthenium-bipyridine-trimethylphosphine caged glutamate (RuBi-glu, Tocris)^68^, diluted into the bath solution for a final concentration of 800 µM. Uncaging of Rubi-glu was performed at 810 nm (∼25-30 mW on sample). The location of the uncaging spot was positioned at ∼ 0.3 µm away from the upper edge of the selected spine head (red dot in Figures 6-8). Changes in calcium were monitored by imaging 2P calcium signals and detecting the fluorescence with 2 PMTs placed after wavelength filters (525/70 for green, 595/50 for red). We performed 2P calcium imaging during five different STDP induction protocols triggered at 0.5Hz: (1) pre-post pairing of +7 ms in one spine; (2) pre-post pairing of +7 ms in two clustered spines; (3) pre-post pairing of +13 ms in one spine; (4) post-pre pairing of −15 ms in one spine; (5) post-pre pairing of −15 ms in two clustered spines. We restricted the image acquisition to a small area (∼150 × 150 pixels) which contained the spine(s) that we uncaged and the shaft. Images were acquired at ∼ 30 Hz, averaged 8 times, with 8 µs dwell time. Calcium signals were imaged 500 ms before STDP induction protocol and right after (4ms) the stimulation for more than 600 ms. We focused our analysis on the images obtained before and immediately after the stimulation in each pairing repetition. ROI drawing was performed using custom algorithms (MATLAB; MathWorks). For spine heads, the ROI was a circle whereas for dendrites it was a polygon. Fluorescence was computed as the mean of all pixels within the ROI. We quantified the relative change in calcium concentration 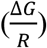 using the following formula:

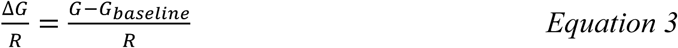

where *G* is the fluorescence from the Fluo-4 dye and *R* is the fluorescence from the Alexa-594 dye. *G_baseline_* is the mean of all pixels of Fluo-4 signal within the ROI taken from the first image at the first stimulation.

### Statistical analysis

Statistics were performed with GraphPad Prism 5. Statistical significance was determined using two-tailed Student’s paired *t*-test when we analyzed the maximum change in uEPSP amplitude after the induction of t-LTP or t-LTD in each experiment and the concomitant changes in the activated spine morphology. Statistical significance was determined using one-way repeated measures ANOVA when we analyzed the time course of the uEPSP amplitude and spine morphological changes after induction of t-LTP or t-LTD with post-hoc pairwise comparisons using Dunnett’s test. *P<0.05; **P<0.01; ***P<0.001.

### Pharmacology

Latrunculin A (Lat-A, Tocris Bioscience) was dissolved in DMSO at 1/1000 and added to the recording chamber containing the brain slice at 100 nM for 15 min before starting the STDP protocol. PEP1-TGL (Tocris Bioscience) was added in the pipette at 200 µM; after 15 min in whole cell condition, electrophysiological recording and synaptic plasticity experiments were started. MNI-glutamate (Tocris Bioscience) was diluted in ACSF from stock solution and bath applied at 2.5 mM. Fresh vials of MNI-glutamate were used for each experiment.

## Ethics

### Animal experimentation

these studies were performed in compliance with experimental protocols (13-185, 15-002, 16-011 and 17-012) approved by the *Comité de déontologie de l’expérimentation sur les animaux* (CDEA) of the University of Montreal.

## ACKNOWLEDGEMENTS

We thank P.J Sjöström and A. Kolta for critical discussion and reading of the manuscript, and are grateful to all other members of Roberto Araya’s laboratory for kind support. We also thank members of the *Groupe de Recherche sur le système nerveux central* (GRSNC) for support and equipment shearing. This work was funded by the Canadian Institutes of Health Research (CIHR) grant MOP-133711 to R.A., a Canada Foundation for Innovation (CFI) equipment grant *Fonds des leaders* 29970 to R.A., and a Natural Sciences and Engineering Research Council of Canada (NSERC Discovery Grant) grant application No. 418113-2012 (NSERC PIN 392027) to R.A. S.T. was supported in part by a salary support from the GRSNC at Université of Montréal. D.E.M. was supported in part by a postdoctoral fellowship from the Fonds de recherche du Québec – Santé (FRQS).

## AUTHOR CONTRIBUTIONS

R.A. conceived the project. S.T. and D.E.M. performed the experiments. S.T. and D.E.M. performed data analyses. S.M-R. performed control experiments. R.A., S.T., and D.E.M. designed experiments. R.A., S.T., and D.E.M. wrote the manuscript. R.A. supervised the project. All authors read and approved the contents of the manuscript.

