## Supplementary Information for "spike-timing-dependent plasticity rule for single, clustered and distributed dendritic spines"

### SUPPLEMENTAL INFORMATION

### Figure S1

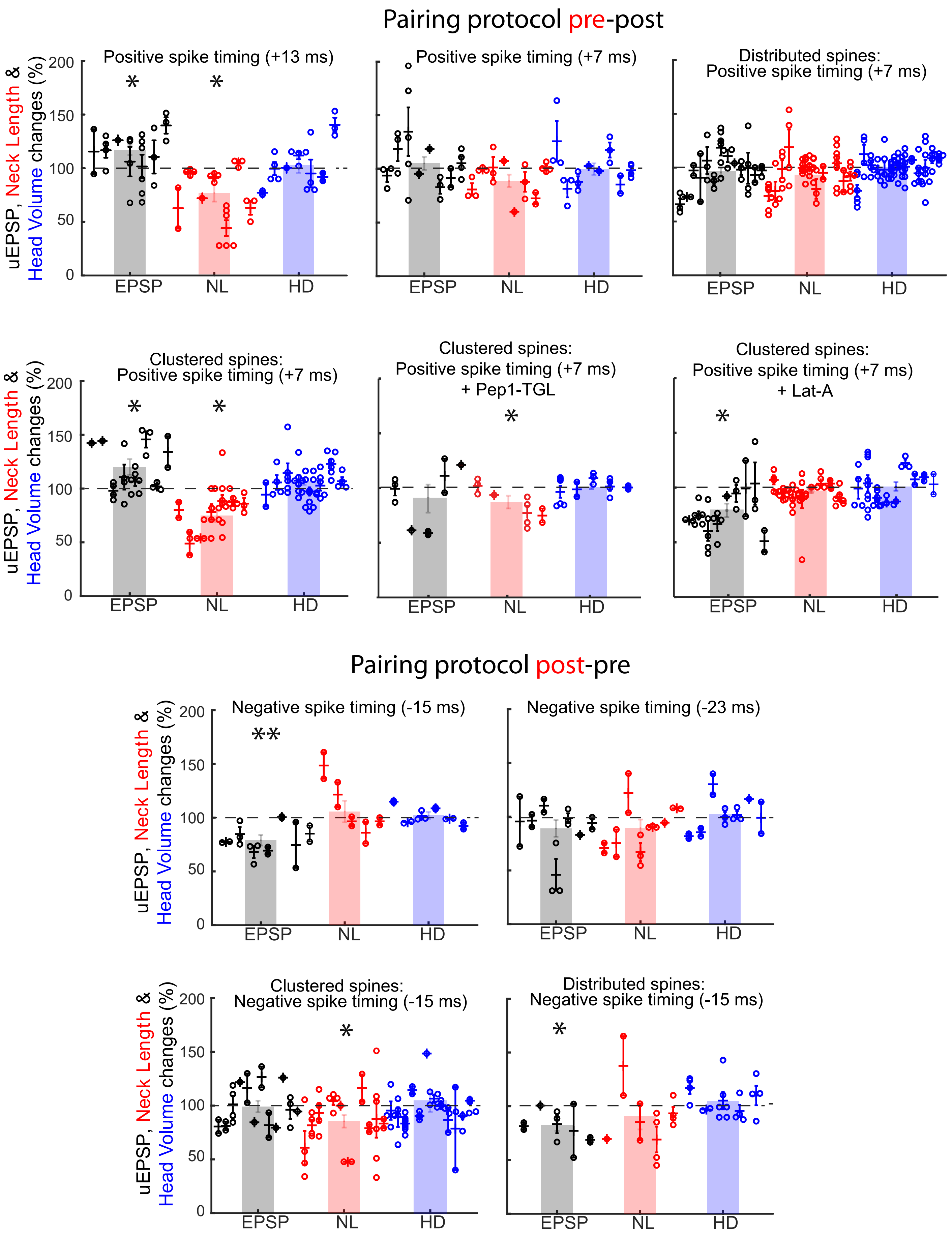

Pairing protocol pre-post
Positive spike timing (+13 ms)
Positive spike timing (+7 ms)
Distributed spines:
Positive spike timing (+7 ms)
Clustered spines:
Positive spike timing (+7 ms)
Clustered spines:
Positive spike timing (+7 ms)
+ Pep1-TGL
Clustered spines:
Positive spike timing (+7 ms)
+ Lat-A
Pairing protocol post-pre
Negative spike timing (-15 ms)
Negative spike timing (-23 ms)
Clustered spines:
Negative spike timing (-15 ms)
Distributed spines:
Negative spike timing (-15 ms)
uEPSP, Neck Length & Head Volume changes (%)
EPSP
NL
HD
0
50
100
150
200
*
**

**Figure S1: Average of all values obtained following STDP induction for uEPSP amplitude, neck length and head volume.** Plots showing all the data points for change in uEPSP (black bars and dots), neck length (red bars and dots) and head volume (blue bars and dots) obtained following STDP induction for each protocol we applied. Each column represents data points from a single experiment. Crosses and errors bars indicate the average and SEM for each individual experiment, while the shaded bar graphs represent the average of the mean from each individual experiment. *$P < 0.05$; **$P < 0.01$; Student's paired-t test. Note that statistical significance remains the same whether we consider the maximum uEPSP change and corresponding changes in morphology (Figure 1-5) or the average of all values obtained following STDP induction for uEPSP amplitude, neck length and head volume.

Figure S2

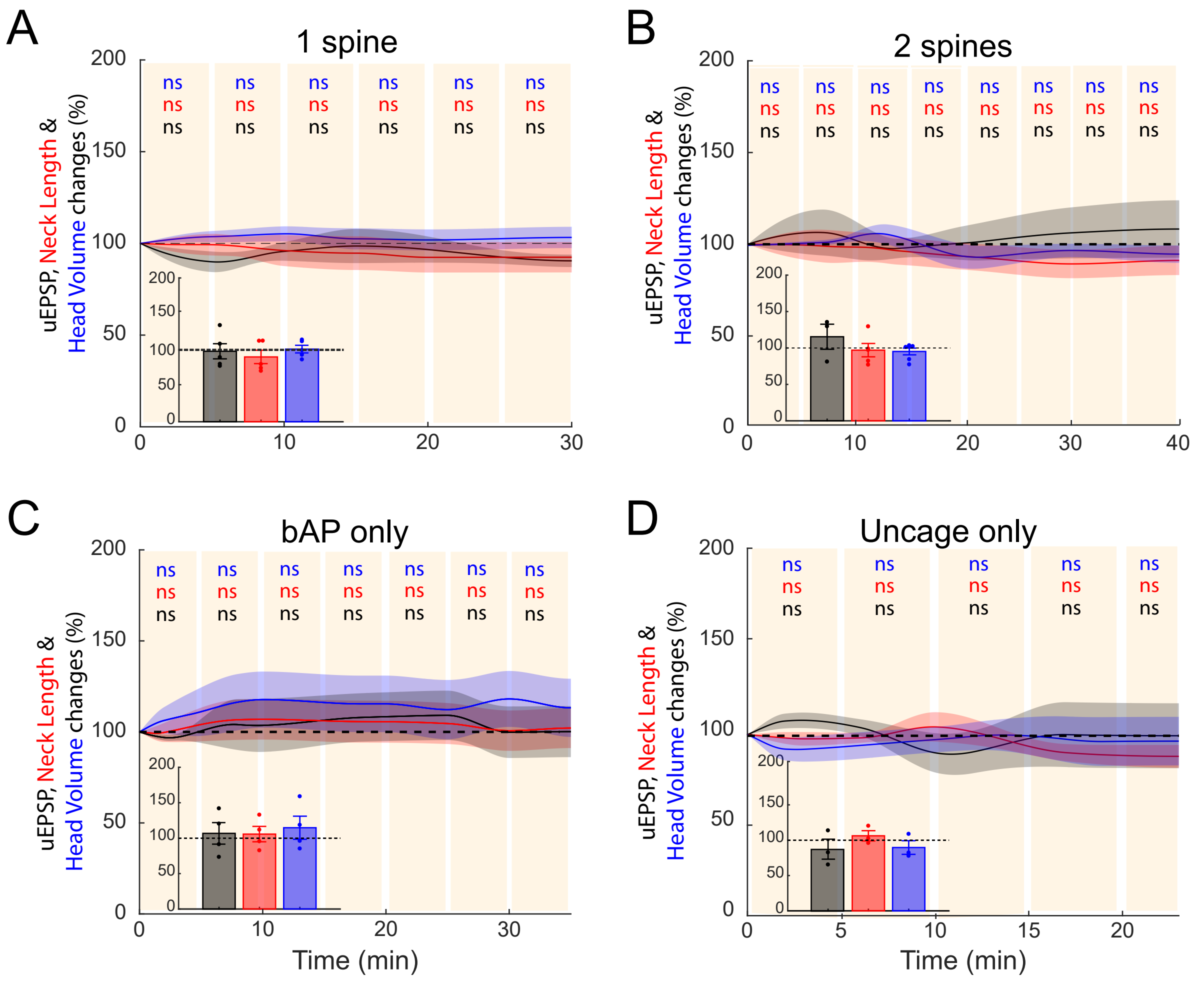

A
1 spine
B
2 spines
C
bAP only
D
Uncage only
uEPSP, Neck Length & Head Volume changes (%)
ns
Time (min)

**Figure S2: Control experiments showing the stability of uEPSP amplitude or spine morphology over time when no pairing protocol was applied.** (A-B) Time course of uEPSP amplitude (black line), neck length (red line) and spine head volume (blue line) over the course of ~30 min without any STDP protocol and uncaging 1 (A) and 2 (B) spines approximately every 5 minutes. Insets show maximum changes in uEPSP amplitude (black bar and dots) and concomitant changes in neck length (red bar and dots) and head volume (blue bar and dots) over the course of ~30 min without any STDP protocol. (C-D) Time course of uEPSP amplitude (black line), neck length (red line) and spine head volume (blue line) over the course of ~30 min following bAP only (C) and synaptic stimulation only (D). Insets show maximum changes in uEPSP amplitude (black bar and dots) and concomitant changes in neck length (red bar and dots) and head volume (blue bar and dots) over the course of ~30 min. *ns*, not significant, one-way repeated measures ANOVA followed by post hoc Dunnet's test.

### Figure S3

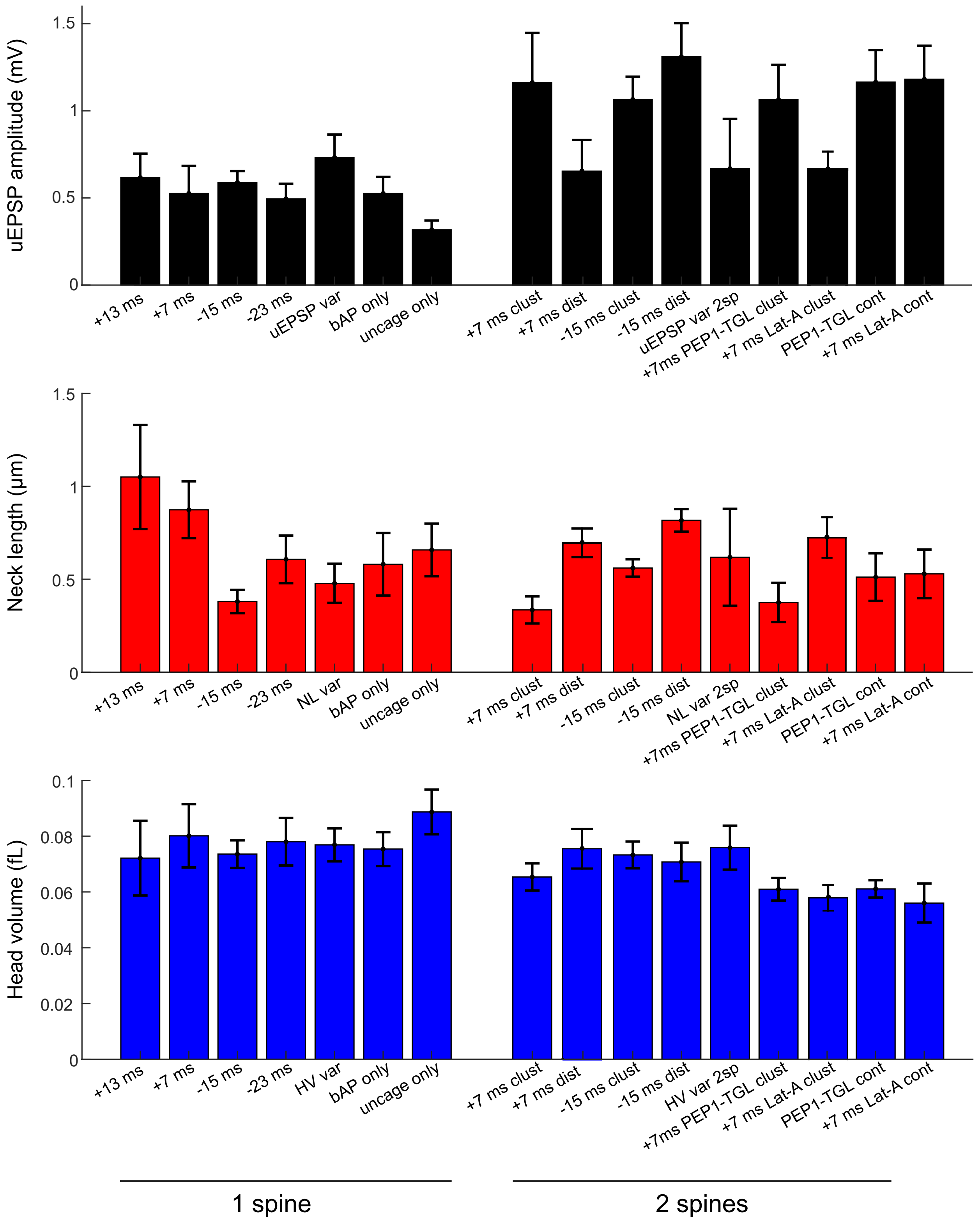

uEPSP amplitude (mV)
1.5
1
0.5
0
+13 ms
+7 ms
-15 ms
-23 ms
uEPSP var
bAP only
uncage only
+7 ms clust
+7 ms dist
-15 ms clust
-15 ms dist
uEPSP var 2sp
+7ms PEP1-TGL clust
+7 ms Lat-A clust
PEP1-TGL cont
+7 ms Lat-A cont
Neck length (μm)
NL var
NL var 2sp
Head volume (fL)
0.1
0.08
0.06
0.04
0.02
HV var
HV var 2sp
1 spine
2 spines

**Figure S3: Actual values for uEPSP amplitude, neck length and head volume.** Bar plots showing the initial values for uEPSP amplitude (black bars), neck length (red bars) and head volume (blue bars) for each STDP protocol applied. A one-way ANOVA followed by a post hoc Tukey's multiple comparison test revealed that the uEPSP amplitude was not significantly different across all conditions when one spine ($P = 0.65$) was activated with two-photon uncaging of glutamate. Similarly, uEPSP amplitude was not significantly different across all conditions when two spines ($P = 0.22$) were activates by 2P uncaging of glutamate. A significant difference in neck length was only found between a pre-post pairing protocol of +13 ms in one spine and +7 ms in two clustered spines ($P < 0.05$; one-way ANOVA followed by a post hoc Tukey's multiple comparison test). The head volume across all conditions was not significantly different ($P = 0.051$; one-way ANOVA followed by a post hoc Tukey's multiple comparison test). uEPSP var, NL var, HV var and uEPSP var 2sp, NL var 2sp, HV var 2sp correspond to the actual values for uEPSP amplitude, neck length, and head volume, respectively, from the experiment shown in Supplementary Figure 2A and B; bAP only, and uncage only correspond to the actual values for uEPSP amplitude, neck length, and head volume from the experiment shown in Supplementary Figure 2C and D, respectively.

Figure S4

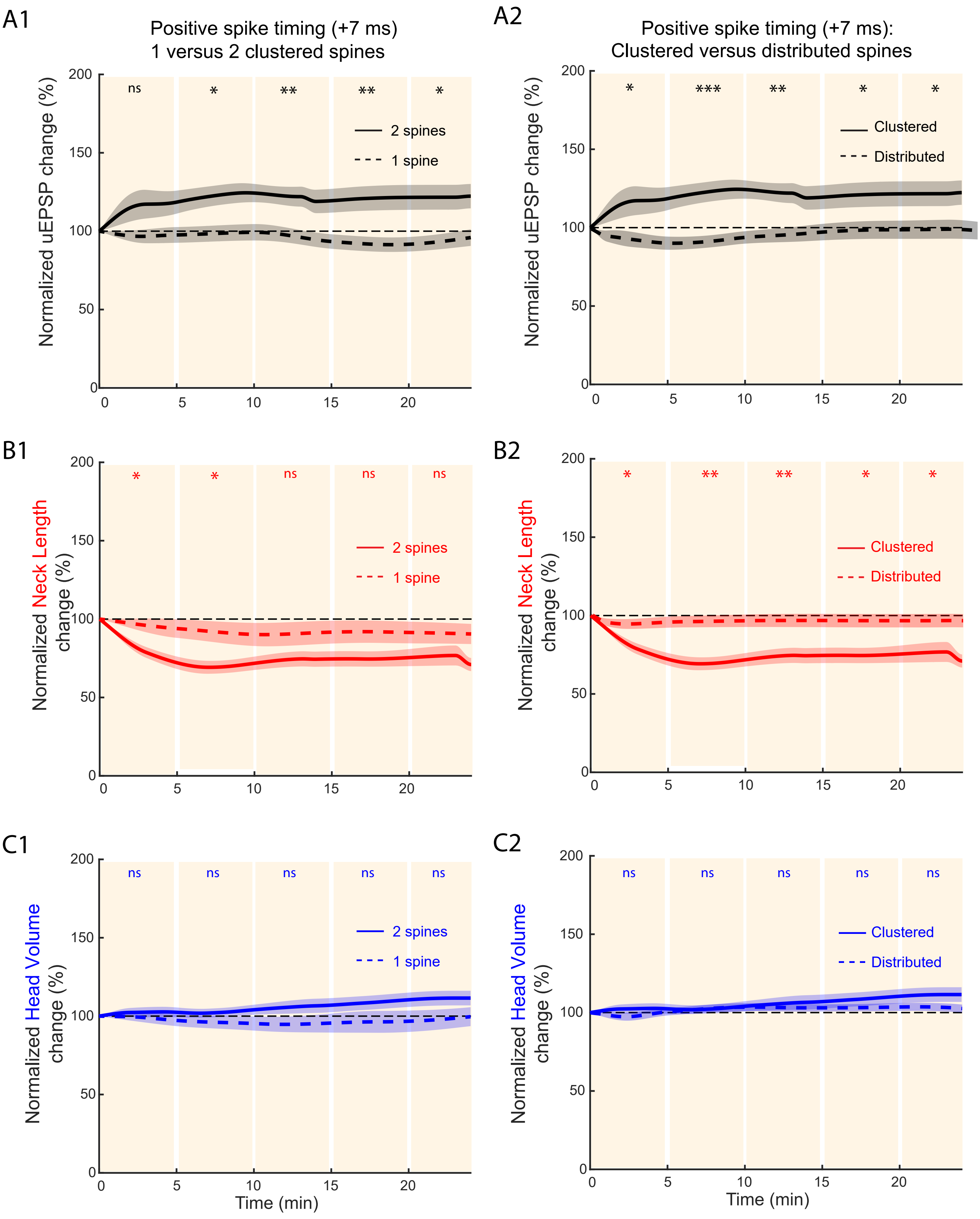
A1
Positive spike timing (+7 ms)
1 versus 2 clustered spines
Normalized uEPSP change (%)
ns
*
**
**
*
2 spines
1 spine
A2
Positive spike timing (+7 ms):
Clustered versus distributed spines
Normalized uEPSP change (%)
*
***
**
*
*
Clustered
Distributed
B1
Normalized Neck Length change (%)
*
*
ns
ns
ns
2 spines
1 spine
B2
Normalized Neck Length change (%)
*
**
**
*
*
Clustered
Distributed
C1
Normalized Head Volume change (%)
ns
ns
ns
ns
ns
2 spines
1 spine
Time (min)
C2
Normalized Head Volume change (%)
ns
ns
ns
ns
ns
Clustered
Distributed
Time (min)

**Figure S4: Induction of t-LTP in single, clustered and distributed dendritic spines at pre-post pairings of + 7ms.** Comparison of the time course of uEPSP amplitude (A), neck length (B) and spine head volume (C) over the course of ~25 min following STDP induction at a pre-post timing of + 7 ms between individual (dashed lines in A.1, B.1 and C1) and two clustered spines (solid lines in A.1, B.1 and C.1), and between two clustered spines (solid lines in A.2, B.2 and C.2) and distributed spines (dashed lines in A.2, B.2 and C.2). *ns*, not significant; $^{*}P < 0.05$; $^{**}P < 0.01$; Student's t-test.

### Figure S5

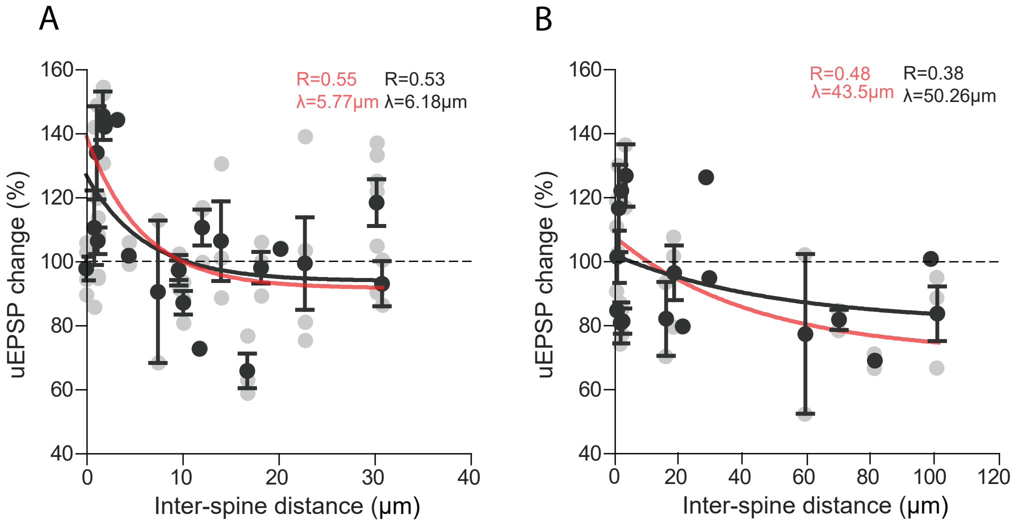

A
R=0.55
λ=5.77µm
R=0.53
λ=6.18µm
uEPSP change (%)
Inter-spine distance (µm)
B
R=0.48
λ=43.5µm
R=0.38
λ=50.26µm
uEPSP change (%)
Inter-spine distance (µm)

**Figure S5: The induction of t-LTP and t-LTD is dependent on the inter-spine distance. (**A) Induction of t-LTP (increase in uEPSP amplitude, more than 100%) in two nearly simultaneously activated spines at a pre-post timing of + 7 ms is dependent on inter-spine distance. Similarly to what was observed when we correlated the inter-spine distance and the maximal change in uEPSP observed in each experiment (Fig. 3E), the mean change (black dots) - obtained when we analyzed the average change in uEPSP from all the times tested in each experiment (gray dots) - following t-LTP induction decayed exponentially as a function of the inter-spine distance with a similar length constant ($\lambda$) ($\lambda$= 5.77 µm and 6.18 µm, respectively, t-test p=0.87). The black line represents an exponential fit to the mean uEPSP change, and the red line represents the fit when the maximum uEPSP change was considered (reported also in Figure 3E). (B) Recovery of t-LTD (decrease in uEPSP amplitude, less than 100%) at a post-pre timing of -15 ms is dependent on inter-spine distance. Similarly to what was observed when we correlated the inter-spine distance and the maximal uEPSP change observed in each experiment (Figure 5D), the mean change (black dots) - obtained when we analyzed the average change in uEPSP from all the times tested in each experiment (gray dots) - following t-LTD induction recovered exponentially as a function of the inter-spine distance with a similar $\lambda$ ($\lambda$= $\lambda$=43.5 µm and 50.26 µm, respectively, t-test, p=0.8). Black line represents the exponential fit to the mean uEPSP change, and red line represents the fit when maximum uEPSP change was considered (reported also in Figure 5D).

Figure S6

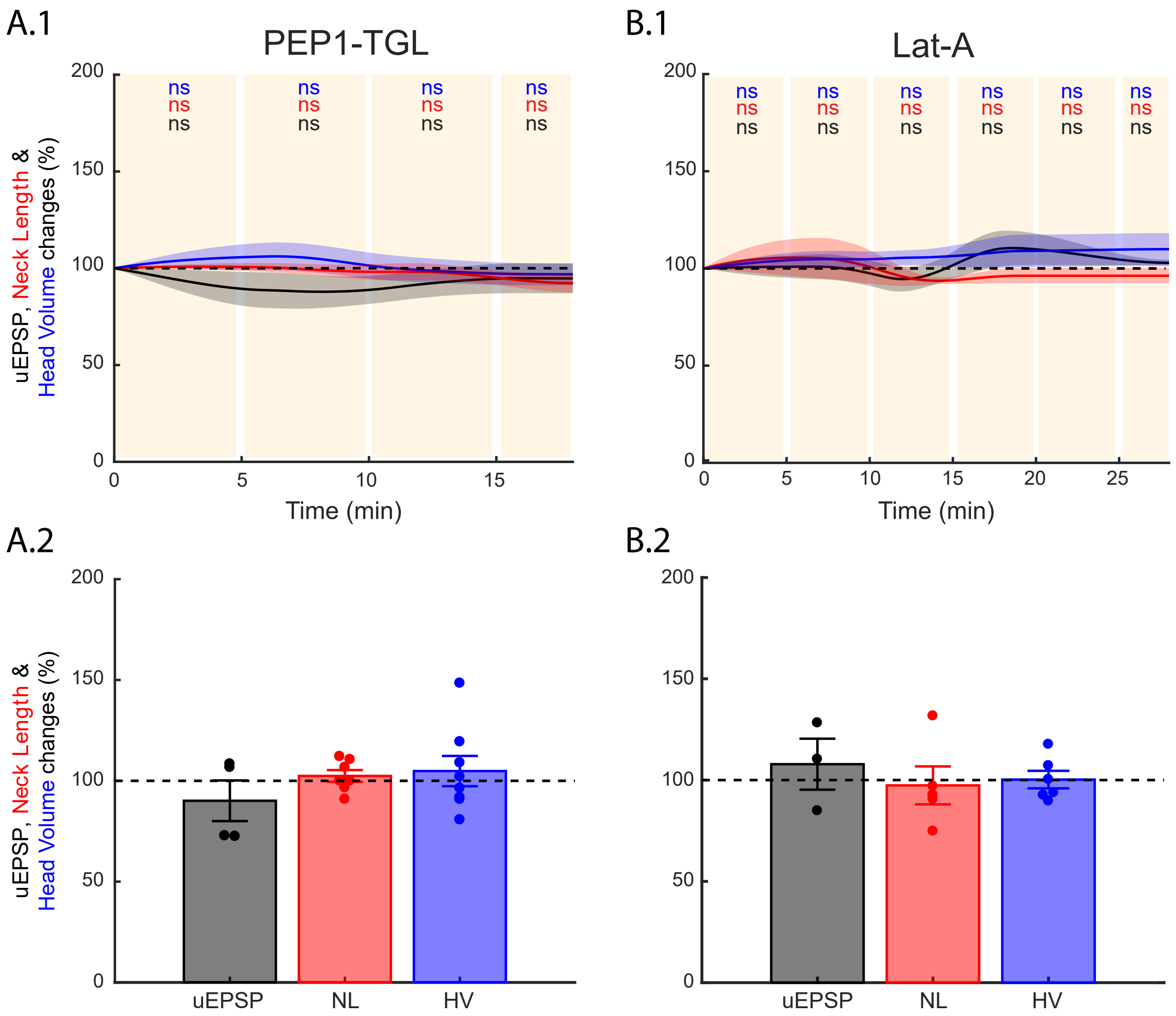

A.1
PEP1-TGL
B.1
Lat-A
ns
ns
ns
uEPSP, Neck Length & Head Volume changes (%)
200
150
100
50
0
0
5
10
15
20
25
Time (min)
A.2
B.2
uEPSP
NL
HV

**Figure S6: Effect of PEP1-TGL and Lat-A on uEPSP amplitude and spine morphology.** Time course of uEPSP amplitude (black line), neck length (red line) and spine head volume (blue line) recorded in the presence of 200 μM PEP1-TGL (A.1) or 100 nM Lat-A (B.1) without any STDP induction protocol. *ns*, not significant, one-way repeated measures ANOVA followed by post hoc Dunnet's test. Maximum changes in uEPSP amplitude (black bar and dots) and concomitant changes in neck length (red bar and dots) and head volume (blue bar and dots) of the activated spine recorded in the presence of 200 μM PEP1-TGL (A.2) or 100 nM Lat-A (B.2) without any STDP induction protocol.

### Figure S7

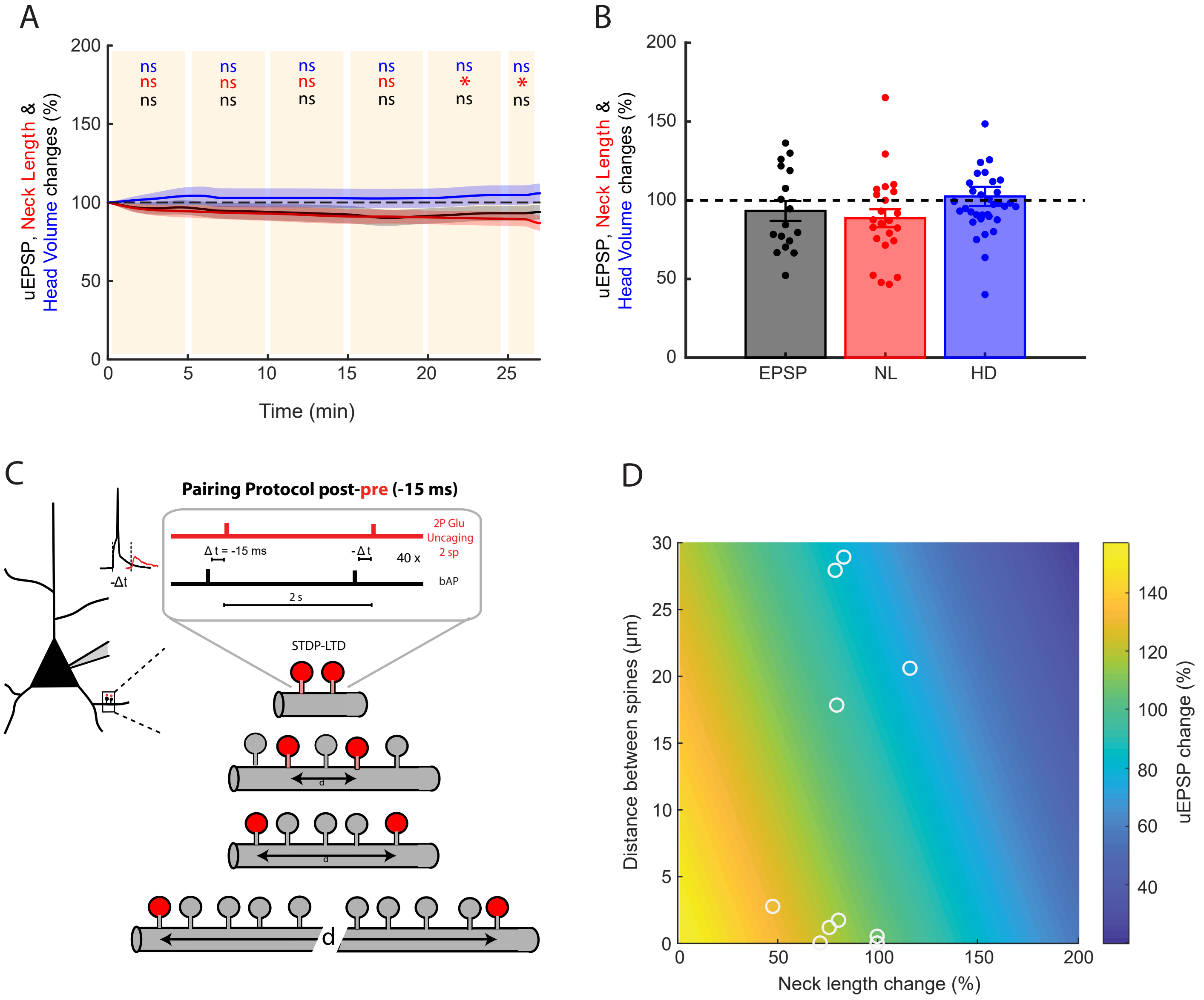

A
B
C
D
uEPSP, Neck Length &
Head Volume changes (%)
ns
*
Time (min)
EPSP
NL
HD
Pairing Protocol post-pre (-15 ms)
2P Glu
Uncaging
2 sp
Δ t = -15 ms
-Δ t
40 x
bAP
2 s
-Δt
STDP-LTD
d
Distance between spines (µm)
Neck length change (%)
uEPSP change (%)

**Figure S7: Recovery of t-LTD following a post-pre pairing protocol of -15 ms in two spines when inter-spine distance is greater than 40 µm.** (A) Time course of uEPSP amplitude (black line), the neck length (red line) and spine head diameter (blue line) of the activated spines for all the inter-spine distances after the induction of t-LTD at pairings of -15 ms. *ns*, not significant; *P < 0.05, one-way repeated measures ANOVA followed by post hoc Dunnet's test. (B) Maximum changes in uEPSP amplitude (black bar and dots) and concomitant changes in neck length (red bar and dots) and head diameter (blue bar and dots) of the two activated spines from each experiment after the induction of t-LTD at a post-pre timing of -15ms. (C) Experimental post-pre induction protocol at pairings of - 15 ms in two dendritic spines separated by different distances. (D) Color plot showing the relationship between uEPSP change (color coded) and neck length change and distance between two clustered spines following a post-pre t-LTD induction protocol. Note that when a pairing protocol of -15 ms is performed in two spines that are close together, and display neck shrinkage, the result is potentiation (increase in uEPSP amplitude, more than 100%). On the other hand, when the induction protocol is performed in two spines that are further away, without neck length changes, the result is depression (decrease in uEPSP amplitude, less than 100%). The change in uEPSP amplitude was modeled using equation 1 (described in methods).

Figure S8

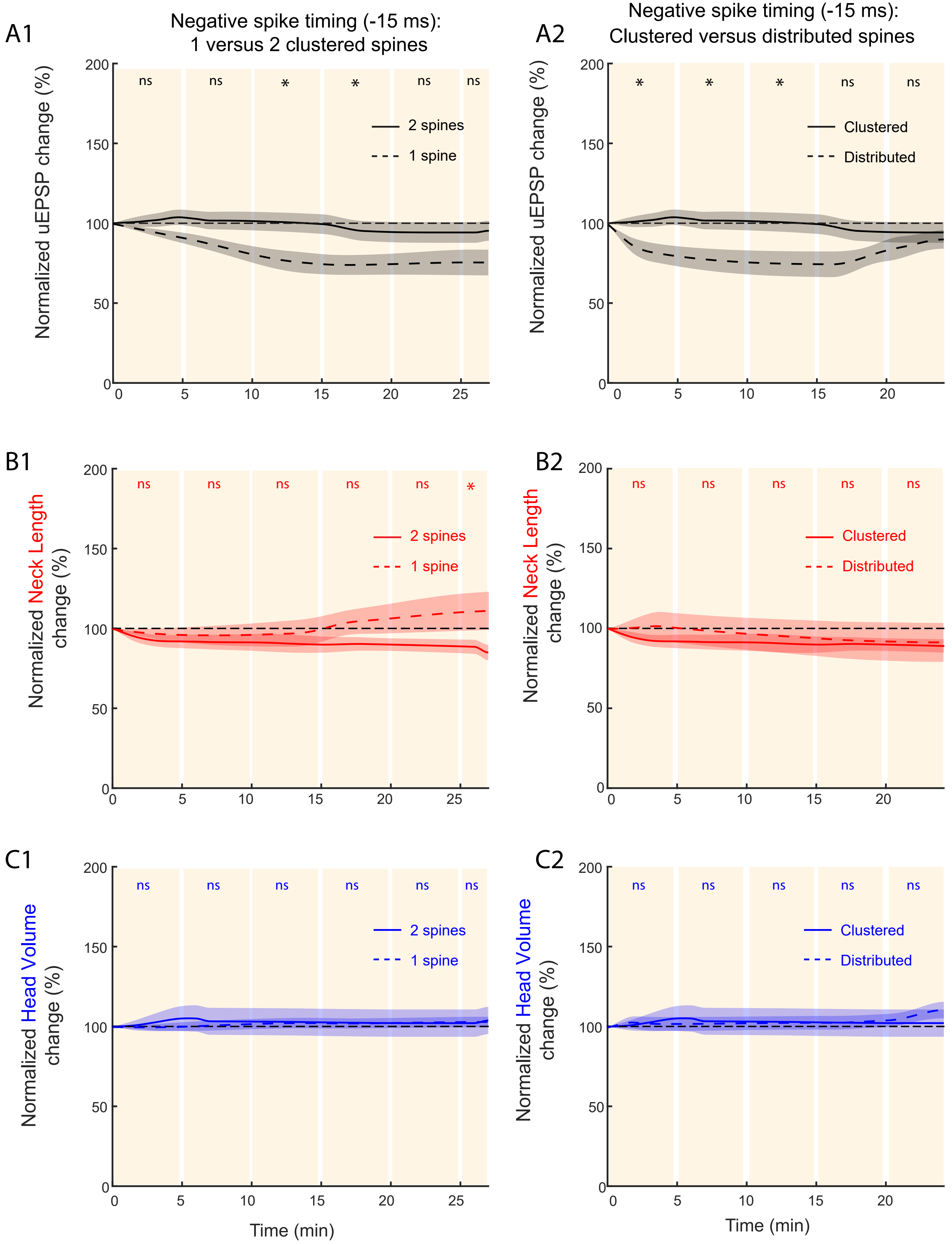

A1
Negative spike timing (-15 ms):
1 versus 2 clustered spines
Normalized uEPSP change (%)
ns
ns
*
*
ns
ns
2 spines
1 spine
A2
Negative spike timing (-15 ms):
Clustered versus distributed spines
Normalized uEPSP change (%)
*
*
*
ns
ns
Clustered
Distributed
B1
Normalized Neck Length change (%)
ns
ns
ns
ns
ns
*
2 spines
1 spine
B2
Normalized Neck Length change (%)
ns
ns
ns
ns
ns
Clustered
Distributed
C1
Normalized Head Volume change (%)
ns
ns
ns
ns
ns
ns
2 spines
1 spine
Time (min)
C2
Normalized Head Volume change (%)
ns
ns
ns
ns
ns
Clustered
Distributed
Time (min)

**Figure S8: Induction of t-LTD in single, clustered and distributed dendritic spines.** Comparison of the time course of uEPSP amplitude (A), neck length (B) and spine head volume (C) over the course of ~25 min following STDP induction at pairings of - 15 ms in individual (dashed lines in A.1, B.1 and C.1) versus two clustered spines (solid lines in A.1, B.1 and C.1), and two clustered spines (solid lines in A.2, B.2 and C.2) versus distributed spines (dashed lines in A.2, B.2 and C.2). *ns*, not significant; *P < 0.05; Student's t-test.

### Figure S9

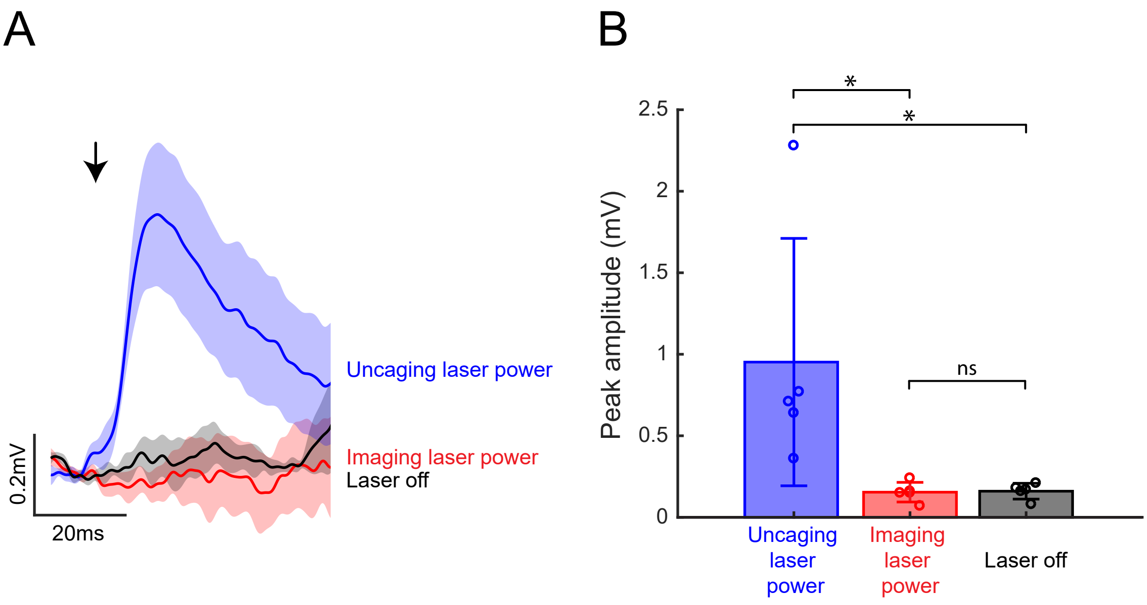
A
0.2mV
20ms
Uncaging laser power
Imaging laser power
Laser off
B
*
*
ns
2.5
2
1.5
1
0.5
0
Peak amplitude (mV)
Uncaging laser power
Imaging laser power
Laser off

**Figure S9: Imaging laser power does not cause glutamate uncaging-mediated uEPSPs in the soma of L5 pyramidal neurons.** (A) Blue trace corresponds to an average of ten depolarizations recorded at the soma caused by uncaging glutamate next to a spine using 4-ms laser pulses of ~25-30mW on sample at 2 second intervals (Uncaging laser power). Note the generation of a clear uEPSP. Red trace corresponds to the average voltage recorded while applying ten 4-ms laser pulses of ~~5 mW on sample at 2 second intervals (Imaging laser power). Note that no uEPSPs were observed. Black trace corresponds to the average voltage recorded a second after the onset of the 4-ms laser pulses, 0mW on sample (Laser off). Shaded area represents the SEM. (B) Plot showing peak amplitude (mV) observed after 2P uncaging of glutamate at Uncaging laser power (Blue), Imaging laser power (red), or with the 2P Laser off (black). N = 5 experiments. *ns*, not significant; *P < 0.05; Student's t-test.

Figure S10

A

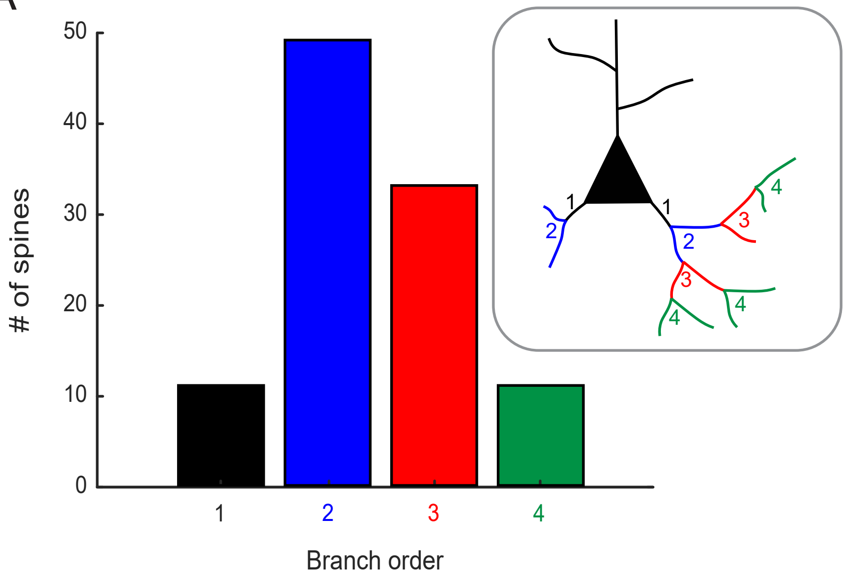


B

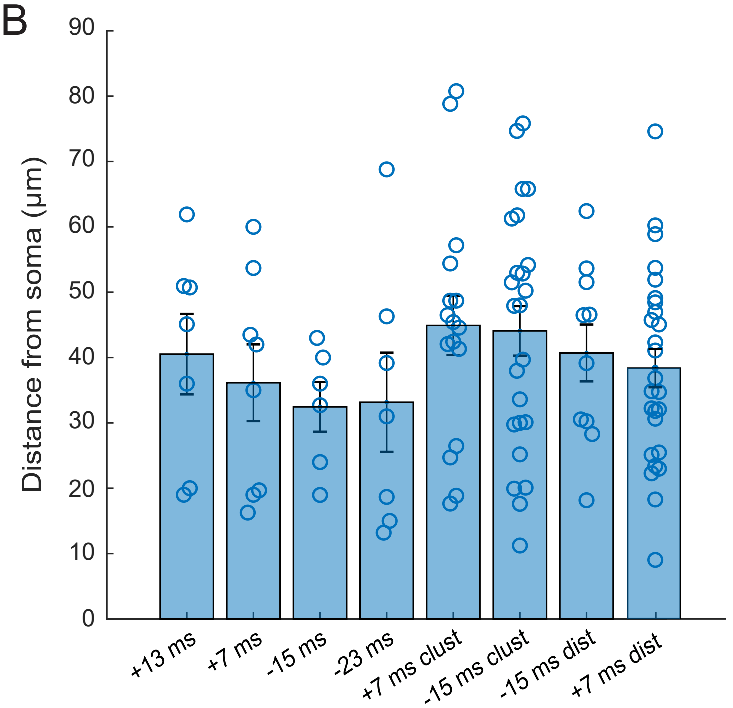

**Figure S10: Morphometric analysis of spines included in this study.** (A) Bar plot showing the branching order of spines that were activated in this study. A primary dendrite is one originating from the cell body (branching order labeled as "1" in inset diagram). The branching order increases with each successive branch point (when dendrite splits into two or more branches). (B) Bar graph showing the distance of spines from the soma for each STDP protocol that we applied. Each data point represents the distance from the soma of individual spines). No significant difference was observed across groups (40.22 ± 1.62 μm away from the soma, $p=0.55$; one-way ANOVA followed by Tukey's Multiple Comparison Test).

Figure S11

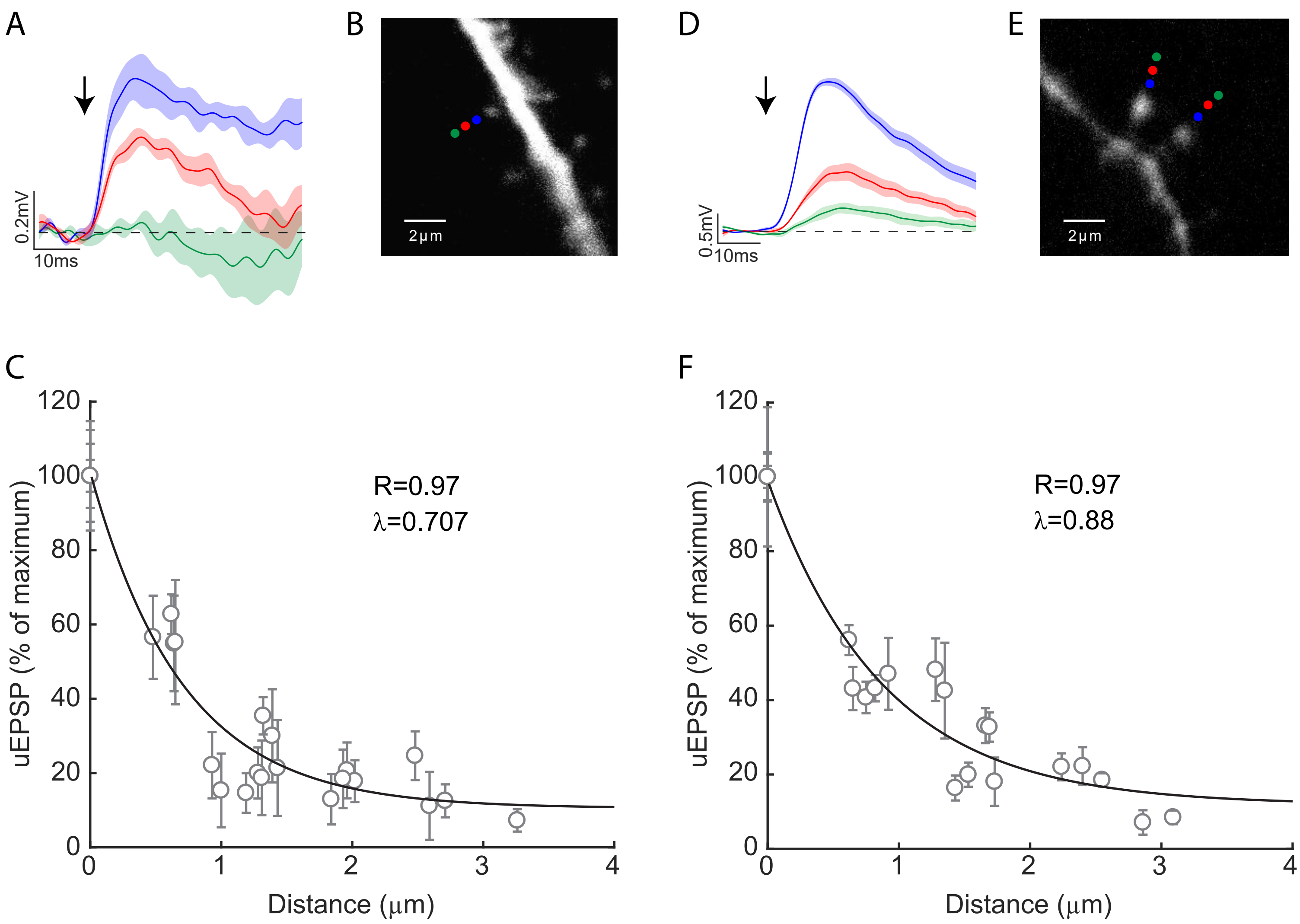

A
0.2mV
10ms
B
2μm
D
0.5mV
10ms
E
2μm
C
uEPSP (% of maximum)
R=0.97
λ=0.707
Distance (μm)
F
uEPSP (% of maximum)
R=0.97
λ=0.88
Distance (μm)

**Figure S11: Spatial resolution of 2P glutamate uncaging in one and two clustered spines.** Two-photon activation of single spines: (A) Example uEPSP averaged traces evoked by placing the uncaging spot at the corresponding color coded locations shown in (B). Each trace corresponds to an average of ten depolarizations recorded at the soma. (C) Averaged uEPSP values (normalized to the maximum value obtained in the same experiment) as a function of distance from the closest uncaging spot in the same experiment. Two-photon activation of two clustered spines: (D) Example uEPSP averaged traces evoked by placing the uncaging spots at the corresponding color coded locations shown in (E). Each trace corresponds to an average of ten depolarizations recorded at the soma. (F) Averaged uEPSP values (normalized to the maximum value obtained in the same experiment) as a function of distance from the closest uncaging spot in the same experiment. Shaded area in A and D represents the SEM.
